# Role of the tomato *MARS1/ROUGH* gene encoding a LYSINE-SPECIFIC HISTONE DEMETHYLASE 1 in adventitious root and fruit skin formation

**DOI:** 10.1101/2025.07.29.665673

**Authors:** Eduardo Larriba, Cécile Bres, Aurora Alaguero-Cordovilla, Johann Petit, Riyazuddin Riyazuddin, Jean Philippe Mauxion, Lara Caballero, Bénédicte Bakan, David Esteve-Bruna, Moussa Benhamed, Christophe Rothan, José Manuel Pérez-Pérez

## Abstract

In contrast to animals, plants have a high regenerative capacity, and they can form new organs and even complete individuals from a few cells present in adult tissues, either in response to injury or to the alteration of their environment. In this study, we describe the isolation and characterization of the *more adventitious roots1-1* (*mars1-1*) mutant, which exhibits enhanced regenerative potential upon wounding in tomato hypocotyl explants. Additionally, the *mars1-1* fruits exhibited a rough surface due to the ectopic proliferation of subepidermal cells, which formed callus-like structures on the cuticle. The *MARS1/ROUGH* gene encodes a conserved lysine-specific histone demethylase, SlLSD1, which regulates a variety of processes in metazoans, including cell proliferation, stem cell pluripotency, and embryogenesis. Two CRISPR/Cas9 null alleles, *mars1-2* and *mars1-3*, were generated and their pleiotropic phenotype was characterized. We found elevated levels of H3K4me1 in *mars1/rough* seedlings, which suggests that SlLSD1 is required for the demethylation of this histone mark. To ascertain the impact of altered epigenetic marks in the *mars1/rough* mutants on gene expression regulation, we conducted a transcriptome analysis using a variety of RNA-Seq studies on tomato hypocotyls. By employing specific bioinformatic workflows and leveraging on the resolution of directional RNA-Seq data, we have identified over several dozen distinct genomic regions that exhibit *de novo* expression in the *mars1/rough* mutants. One such region includes a novel B-type cyclin gene, which is upregulated in the *mars1/rough* mutants and may account for the observed phenotypes. Our findings indicate that SlLSD1 plays a role in the establishment and maintenance of silencing in specific genomic regions that are essential for tissue-specific reprogramming.

## Introduction

Plants display an impressive capacity for regeneration, enabling them to repair damaged tissues and even regenerate lost organs (Xu and Huang, 2014; Ikeuchi et al., 2016; Ikeuchi et al., 2019). In the context of environmental stress, the regeneration of damaged plant parts has been demonstrated to be a consequence of hormonal crosstalk, thereby restoring normal tissue function. Following injury, various tissues possess the capacity to induce regeneration of missing organs, resulting in the formation of adventitious roots (ARs) or shoots. The results of studies conducted on the model plant *Arabidopsis thaliana* have identified several transcription factors and hormonal crosstalk as key players in wound-induced organ regeneration (Mathew and Prasad, 2021).

The tomato cultivar ‘Micro-Tom’ (MT) serves as an excellent model for research in Solanaceae due to its similarities with Arabidopsis, including its small size, short life cycle, and simplicity of transformation (Shikata and Ezura, 2016). The implementation of an optimized CRISPR-Cas9 editing system enables the utilization of reverse genetics methodologies (Dahan-Meir et al., 2018). In previous studies (Alaguero-Cordovilla et al., 2021; Larriba et al., 2021a; Larriba et al., 2021b; Larriba et al., 2022), we established a new experimental system to study regeneration in tomato, using hypocotyl explants of MT to simultaneously study wound-induced *de novo* shoot formation and AR development without exogenous application of hormones. In our working model, the cellular reprogramming of the founder cells of the new organs requires a very precise spatial and temporal regulation of auxin and cytokinin (CK) gradients in the most apical and basal regions of the explants, coupled with an extensive metabolic and dynamic homeostasis of reactive oxygen species (ROS). We also identified several conserved transcription factor modules, such as those homologs of WOUND-INDUCED DEDIFFERENTIATION 1 - ENHANCER OF SHOOT REGENERATION 1/2 - WUSCHEL-RELATED HOMEOBOX 13 (WIND1-ESR1/2-WOX13) and SOLITARY ROOT/INDOLE-3-ACETIC ACID14 -AUXIN RESPONSE FACTOR 7/19 - LATERAL ORGAN BOUNDARIES-DOMAIN (SLR/IAA14-ARF7/ARF19-LBD), involved in the specification of callus and root identity during *de novo* shoot and AR formation after wounding, respectively. We also found a significant variation in the expression of long non-coding RNAs (lncRNAs), both depending on the regions studied and over time (Larriba et al., 2021a), suggesting a regulatory role of these non-coding sequences in wound-induced organ formation in tomato.

Recent work in tomato hypocotyl explants (Yang et al., 2024) has characterized loss-of-function mutants of *REGENERATION FACTOR 1* (*REF1*), which exhibited reduced callus formation and shoot regeneration upon exogenous hormone treatment. In contrast, plants overexpressing *REF1* showed significantly increased regeneration capacity. REF1 encodes a small peptide that induces the expression of *SlWIND1*, the tomato homolog of *WIND1* in Arabidopsis that promotes wound-induced callus formation in many species (Iwase et al., 2017).

To gain some insight into the genetic mechanisms that modulate wound-induced AR formation in tomato explants, we have screened a highly-mutagenized ethyl methanesulfonate (EMS) mutant population generated in the miniature cultivar MT at INRA Bordeaux (Just et al., 2013). Most of the AR mutants identified in our screen displayed some defect in AR initiation, which causes a complete absence or a reduced number of ARs. We have initiated in this work the molecular characterization of mutants with an increased number of ARs, which might be defective in some of the negative regulators of wound-induced AR formation, and that we dubbed as *more adventitious roots* (*mars*) mutants. One of these mutants, *mars1/rough*, has also been independently identified in a genetic screen for tomato mutants affected by fruit cuticle cracking. The present study reports the results of a genetic and molecular analysis of these mutants. Our findings indicate that *MARS1/ROUGH*, which encodes a conserved lysine-specific histone demethylase, plays a key role in a novel genomic surveillance mechanism to broadly downregulate silenced regions within euchromatin.

## Results

### Identification of Mutants Affected in Wound-Induced Adventitious Root Formation and Fruit Skin Formation

A highly mutagenized collection of lines generated by EMS in *S. lycopersicum* cv. MT was evaluated for early seedling and root system architecture (RSA) mutant phenotypes. A small number of the lines exhibited seedlings with an increased number of ARs following the excision of the entire root system (Alaguero-Cordovilla et al., 2020). These may have been the consequence of mutations that affected the negative regulators of wound-induced AR formation. The P19E6 M_3_ line showed a segregation pattern for mutant seedlings, characterized by a significant increase in wound-induced ARs compared to its MT siblings (**Fig. 1A, B**). The observed phenotypic proportions were consistent with the recessive inheritance of the mutant phenotype. The primary root (PR) length was significantly shorter in the P19E6 mutants compared to the MT background (**Fig. 1C**), mainly due to the slight delay in germination observed in the mutants (**Suppl. Fig. S1A**). The root hairs in the P19E6 mutants were closer to each other (**Fig. 1D**) though there were no significant differences in their length compared to the MT siblings (**Suppl. Fig. S1B**). Subsequently, the number of lateral roots (LRs) was quantified following the surgical excision of the PR tip, as previously described (Alaguero-Cordovilla et al., 2020). No significant differences were observed between the P19E6 mutants and MT in the number or density of LRs (**Suppl. Fig. S1C-E**).

**Fig. 1.**
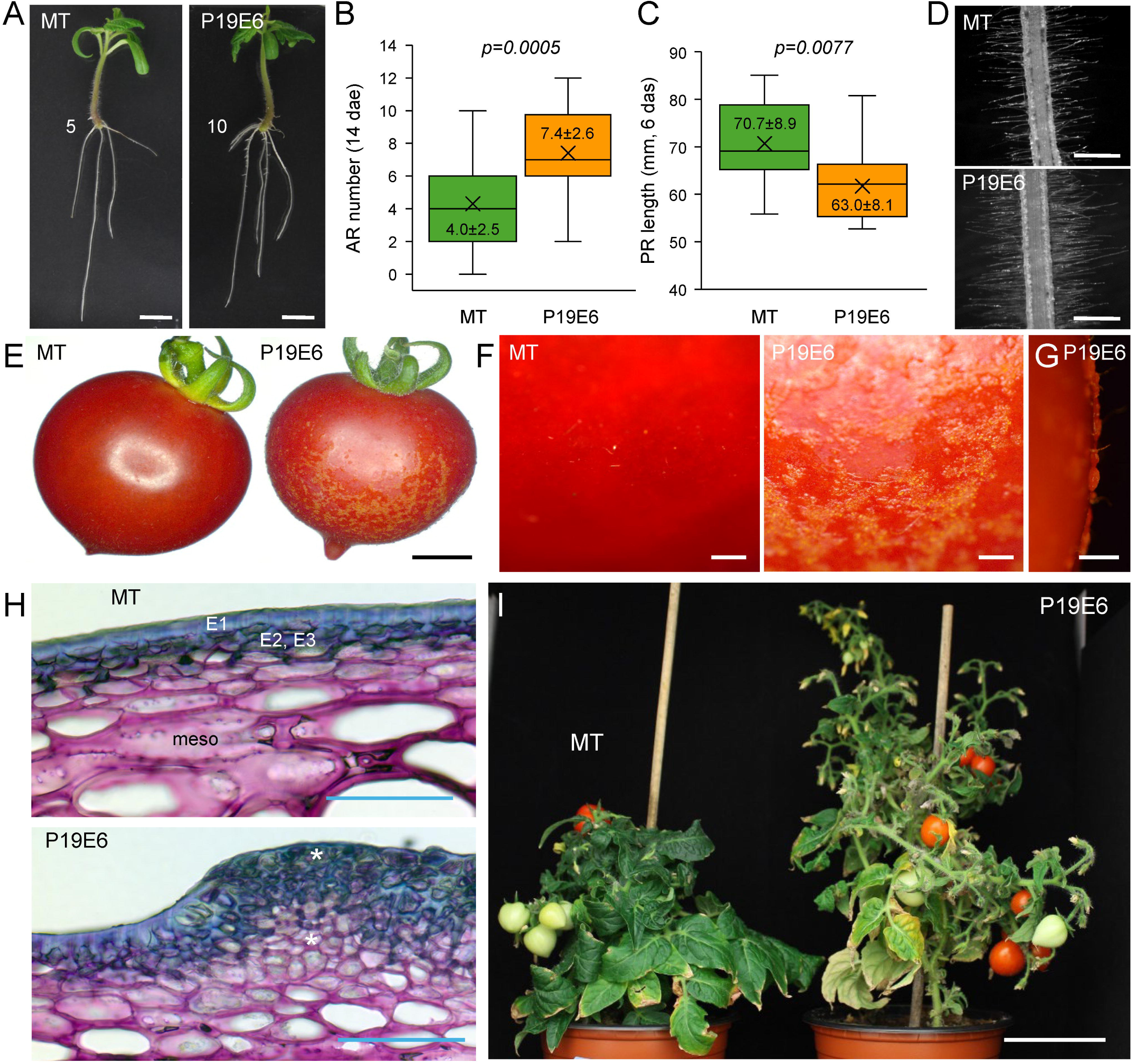
Morphological phenotype in the P19E6 mutant. (A) Representative images of rooted shoot explants at 14 days after excision (dae). The number of ARs is indicated. (B-C) Box plot representation of AR numbers (B) and PR length (C). The numbers indicate the mean ± standard deviation. The p-values are indicated, with those in italics indicating statistical significance (T-test; p-value<0.05; n>18). (D) Representative images of the differentiated region of the PRs, which exhibited fully developed root hairs. (E to G) Macroscopic observations of the fruit surface in MT and the P19E6 mutant at the red ripe stage. (H) Toluidine blue staining of the outer pericarp section of the mature green stage of the fruit in MT and P19E6 fruits. Legend: E1, outer epidermal layer of exocarp; E2 and E3 layers of exocarp, meso, mesocarp layers. The asterisks indicate the presence of meristematic cells in the region of the outer pericarp where the knobs are found in the P19E6 mutant. (I) Representative images of the vegetative phenotype of adult plants after 7 weeks in the growth chamber. Scale bars: 5 cm (I), 10 mm (A, E), 500 µm (F, G), and 150 µm (C, H).

The P19E6 mutant was also independently identified in a genetic screen for tomato mutants affected by fruit cuticle cracking. At the red ripe stage, this mutant exhibited cuticular defects on the fruit surface including skin microcracks compared to MT fruits (**Fig. 1E**). Detailed observation of the surface of P19E6 mutant fruits further revealed that lighter-colored patches of tissue protruded outwards from the fruit skin (**Fig. 1F**), which resulted in a rough surface in these areas (**Fig. 1G**). Histological study of cross sections of fruits in the P19E6 mutants confirmed the presence of supernumerary cells in the exocarp, leading to outgrowth of the fruit skin (**Fig. 1H**). Uncontrolled outward cell proliferation was initiated as early as 5 days post anthesis (DPA; **Suppl. Fig. S1F**). The analysis of the fruit cuticle of the P19E6 mutants showed an almost 3-fold increase of the cutin polymer load (3.29±0.23 mg/cm^2^ in P19E6 and 1.23±0.61 mg/cm^2^ in MT fruits; *p*-value = 0.005), whose composition was reminiscent of that of suberin, displaying an increased abundance of very long chain fatty acids such as 20-OH-C20:0 (**Suppl. Fig. S1G**). The wax load increased by almost 2-fold, but its composition remained unaffected. The P19E6 mutants also showed an altered floral phenotype, characterized by more branching and more flower trusses than MT (**Fig. 1I**). Based on the AR and fruit skin phenotypes, the P19E6 mutants will be herein referred to as the *more adventitious roots1-1* (*mars1-1*) or *rough-1*.

### Molecular Identification of *MARS1/ROUGH* Gene Encoding a LYSINE-SPECIFIC HISTONE DEMETHYLASE 1 Protein

Segregation analysis of the rough fruit phenotype in the BC_1_F_2_ population issued from a P19E6 × MT cross confirmed the presence of a single recessive mutation. To identify the mutation responsible for this phenotype, we used an established mapping-by-sequencing strategy (Petit et al., 2016; Garcia et al., 2016). The whole-genome sequencing of a bulk of 44 BC_1_F_2_ plants displaying the rough fruit phenotype followed by the analysis of the allelic frequency distribution of EMS-induced mutations revealed a dramatic increase in mutation frequencies on chromosome 10 (**Fig. 2A**). Subsequent fine mapping using 768 BC_1_F_2_ recombinant plants allowed us to unequivocally identify the causal mutation of the rough fruit phenotype, which is a G to A transition in the coding region of *Solyc10g047350* (**Fig. 2B**). This gene encodes a protein with homology to the LYSINE-SPECIFIC HISTONE DEMETHYLASE 1 (LSD1), also known as lysine (K)-specific demethylase 1A (KDM1A) (Shi et al., 2004) (**Fig. 2D**). The P19E6 mutation changes a glycine (G) to an arginine (R) residue at position 227 of the SlLSD1 protein, within the SWIRM (<u>Sw</u>i3p, <u>R</u>sc8p and <u>M</u>oira) domain, which is found in many chromosomal proteins involved in chromatin modification (Da et al., 2006). This mutation affects a highly conserved glycine residue in all tomato and Arabidopsis proteins with homology to LSD1/KDM1A (**Fig. 2E**; Gao et al., 2012), which produces a evident alteration of the 3D structure in the SWIRM domain (**Fig. 2F**). LSD1/KDM1A also contains the amine oxidase domain, which is responsible to catalyze the oxidation of methylated lysines in the histone H3 tail (Shi et al., 2004).

**Fig. 2.**
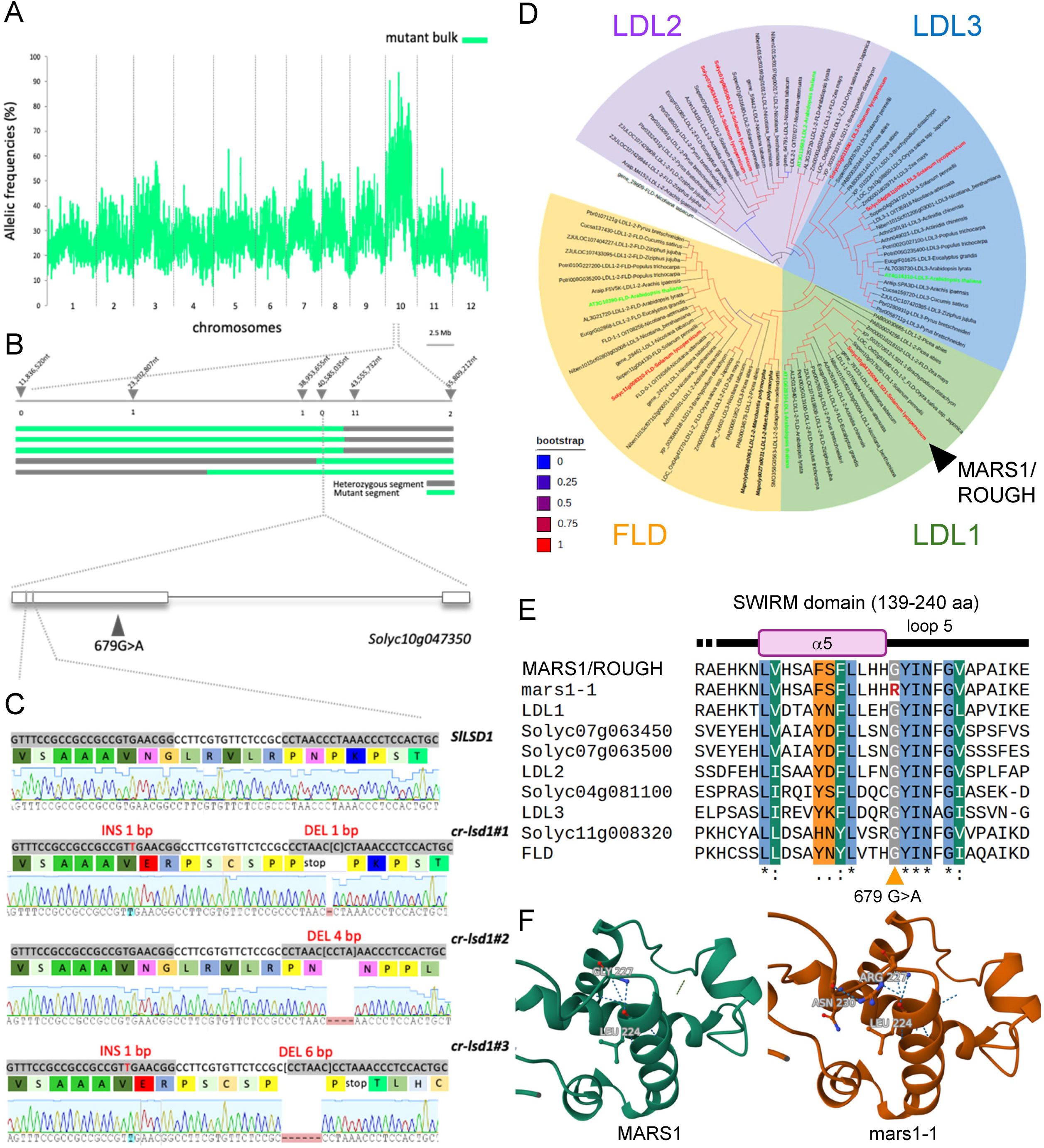
Identification of the gene responsible for the pleiotropic *mars1/rough* phenotype. (A) Mapping-by-sequencing of the P19E6 mutant. Pattern of mutation allelic frequencies obtained in the mutant bulks is represented along tomato chromosomes by a green line. Chromosome 10 exhibits a dramatic increase in mutation allelic frequency. (B) Fine mapping of the causal mutation using BC_1_F_2_ population. Recombinant scoring of 768 BC_1_F_2_ individual plants allows accurate identification of the causal mutation at position 40,585,035 on chromosome 10 (SL2.50, ITAG2.4). The P19E6 EMS mutant contains a G-to-A transversion at position 679 of the *Solyc10g047350* gene, encoding a protein homologous to lysine-specific histone demethylase 1 (LSD1). (C) CRISPR/Cas9-induced mutations in the first exon of *SlLSD1* obtained in three independent homozygous lines. Bases highlighted in grey indicate the two guide RNA target sequences. Insertions (INS) or deletions (DEL) are indicated in red. CRISPR/Cas9-induced frameshift insertions and deletions lead to truncated SlLSD1 proteins. (D) Phylogeny of full-length LSD1-like homologs in representative plant species. Green and red, respectively indicates *A. thaliana* and *S. lycopersicum* sequences. (E) Partial alignment of the SWIRM domain of the LSD1-like family proteins from *A. thaliana* (LDL1, LDL2, LDL3, and FLD) and *S. lycopersicum* (SlLSD1). The *mars1-1* mutation present in the P19E6 line resulted in the replacement of a highly conserved glutamine with arginine. (F) 3D structure of the protein region surrounding the mutation found in the P19E6 (*mars1-1*) line, according to the models predicted by AlphaFold (Jumper et al., 2021).

We next generated several gene-edited CRISPR/Cas9 lines in MT and selected three of these lines showing different mutations (**Fig. 2C** and **Suppl. Table S1**). Line *mars1-2* carries a one-base insertion between positions 287 and 288 and a one-base deletion at position 315 from the ATG, which generates a frameshift and a truncated protein of 104 amino acids due to the appearance of a premature termination codon (**Fig. 2C**). Line *mars1-3* presents the insertion of a base pair between positions 287 and 288 and the deletion of six base pairs at position 310-315 from the ATG, resulting in the production of a truncated protein of 104 amino acids due to the appearance of a premature termination codon after the frameshift (**Fig. 2C**). Gene editing of line *mars1-4* resulted in a four-base pair deletion at position 316-319 from the ATG, which generates a truncated protein of 118 amino acids due to the appearance of a premature termination codon after the frameshift (**Fig. 2C**). For further studies, we chose the gene-edited lines *mars1-2* and *cr-lsd1#3*, which exhibited distinct mutations and yet produced analogous mutant proteins. These proteins were comprised solely of a short amino-terminal region of the protein (104 amino acids), which was non-functional due to the absence of the SWIRM and amino oxidase domains.

The *mars1-2* and *mars1-3* mutants exhibited increased AR production in shoot explants following whole root excision, compared to the MT background, with more pronounced phenotypes than those observed in the *mars1-1* mutant (**Fig. 3A, B**). In addition, both *mars1-2* and *mars1-3* showed the rough fruit phenotype of *mars1-1,* this phenotypic alteration being more pronounced in *mars1-3*, where it was concomitant with increased branching and flower production (**Fig. 3F** and **Suppl Fig. S2A-C**). A detailed microscopic examination of the developing fruit exocarp revealed that the *mars1-2* and *mars1-3* mutants exhibited aberrant periclinal divisions and subsequent cell proliferation in the fruit epidermis during the cell division phase (5-8 DPA). This resulted in the formation of disorganized epidermal cell masses on the fruit surface (**Fig. 3F, G**). During the cell expansion phase (20 DPA), the loss of cell adhesion in these callus-like structures was followed by the rupture of the fruit skin in protruding areas (**Fig. 3G**). During the phenotypic characterization of the CRISPR/Cas9 mutants of *SlLSD1*, it was observed that a significant proportion of the seeds of the mutants did not germinate or exhibited early lethality (**Fig. 3H, I**). These results are consistent with the more severe molecular damage observed in the two CRISPR/Cas9 mutants than in the EMS-induced *mars1-1* mutants. This suggests that *mars1-2* and *mars1-3* mutants might carry null mutations in the *SlLSD1* gene, while *mars1-1* could be a hypomorphic mutation resulting in a mild loss of function in *SlLSD1*.

**Fig. 3.**
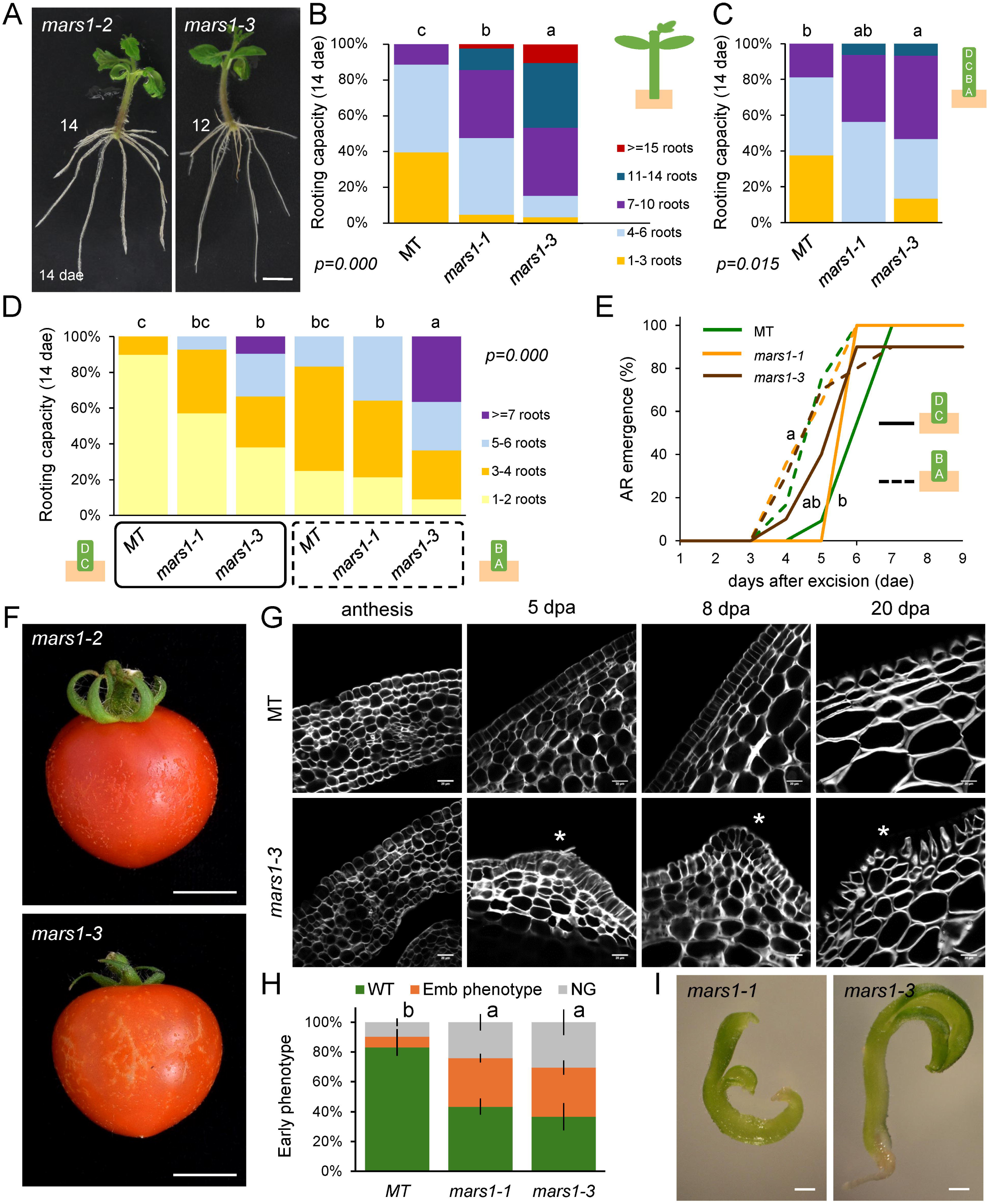
AR formation and fruit phenotype in *mars1/rough* mutants. (A) Representative images of rooted shoot explants at 14 days after excision (dae). The number of ARs is indicated. (B) Rooting capacity of shoot explants at 14 dae. (C, D) Rooting capacity of hypocotyl explants at 14 dae. (E) AR emergence in the studied explants. Different letters indicate significant differences between the studied conditions (LSD; p-value<0.05; n>15). (F) Macroscopic observations of the fruit surface in *mars1/rough* mutants at the red ripe stage. (G) Calcofluor staining and confocal scanning microscopic images of pericarp tissue cells of MT and *mars1-3* fruits; dpa, day post anthesis. The asterisks indicate the presence of meristematic cells in the region of the outer pericarp where the knobs are found in the *mars1/rough* mutants. (H) Early seedling genotype. Emb: seedlings resembling mutants defective in embryo development, NG: not germinated seeds. Letters indicate significant differences (LSD; p-value<0.05; n>225) between genotypes. (I) Representative images of Emb phenotypes in *mars1/rough* mutants at 10 days after sowing. Scale bars: 1 mm (I), 10 mm (A), and 20 µm (G).

### *mars1/rough* Mutants are Affected in the Auxin-to-Cytokinin Responses that Regulate *De Novo* Organ Formation

Previously, we characterized wound-induced AR formation in young shoot and hypocotyl explants in several tomato cultivars, including MT (Alaguero-Cordovilla et al., 2021). Our findings indicated that basipetal auxin transport from the shoot and local auxin biosynthesis triggered by wounding play a pivotal role in this process (Alaguero-Cordovilla et al., 2021). We found that the *mars1/rough* mutations also enhanced the rooting capacity of whole hypocotyl explants without the shoot (**Fig. 3C**), as well as that of smaller explants obtained from sectioning hypocotyls in halves, corresponding to the apical or basal region of the hypocotyl (**Fig. 3D**). The emergence of ARs was observed to be significantly delayed in apical-derived explants compared to those of basal-derived explants, regardless of genotype (**Fig. 3E**). Transverse sections of the basal region of the hypocotyl revealed that the AR primordia arise from proliferation of the interfascicular cambial cells that are on the side facing the phloem poles and adjacent to the vascular bundle, in both the MT background (**Fig. 4A, A’**) and *mars1/rough* mutants (**Fig. 4B, B’**). The observed increase in the number of ARs in the *mars1* mutants cannot be attributed to the longitudinal expansion of the AR formative region at the base of the hypocotyl. Rather, it appears to be the result of an increase in the radial density of ARs, as evidenced by cross sections (**Fig. 4C, D**). These results are consistent with the hypothesis that *mars1/rough* mutations could have altered the responses to the dynamic hormonal balance between auxins and CKs, which is necessary for the specification of new AR primordia from the cambium (Larriba et al., 2021b). The dual-color transcriptional reporter *DR5:mScarletI-NLS*;*TCSn:mNeonGreen-NLS* was employed to study endogenous auxin and CK responses (Omary et al., 2022). At the time of wounding (0 h after excision, hae), the expression of the CK response marker *TCSn:mNeonGreen-NLS* was observed in numerous cells of the vascular cylinder in the MT hypocotyls. In contrast, the expression of the auxin response marker *DR5:mScarletI-NLS* was found to be significantly lower in these cells, with some overlap between the two markers being expressed in specific cells, as evidenced by double blue (CK response) and red (auxin response) fluorescence (**Fig. 4C’**). Similar results were observed in the *mars1-1* mutants (**Fig. 4D’** and **Suppl. Fig. S3**). At 40 and 75 hae, the expression of the CK reporter (TCS) in the vascular stele significantly decreased in both MT and *mars1-1* mutants (**Fig. 4E** and **Suppl. Fig. S3**). The most conspicuous differences between MT and *mars1-1* were due to the enhanced signal shown by the auxin reporter (DR5) in the earlier time point in the latter (**Fig. 4E** and **Suppl. Fig. S3**). These results indicate that the *mars1-1* mutants exhibited increased auxin responsiveness after wounding when compared to the MT background.

**Fig. 4.**
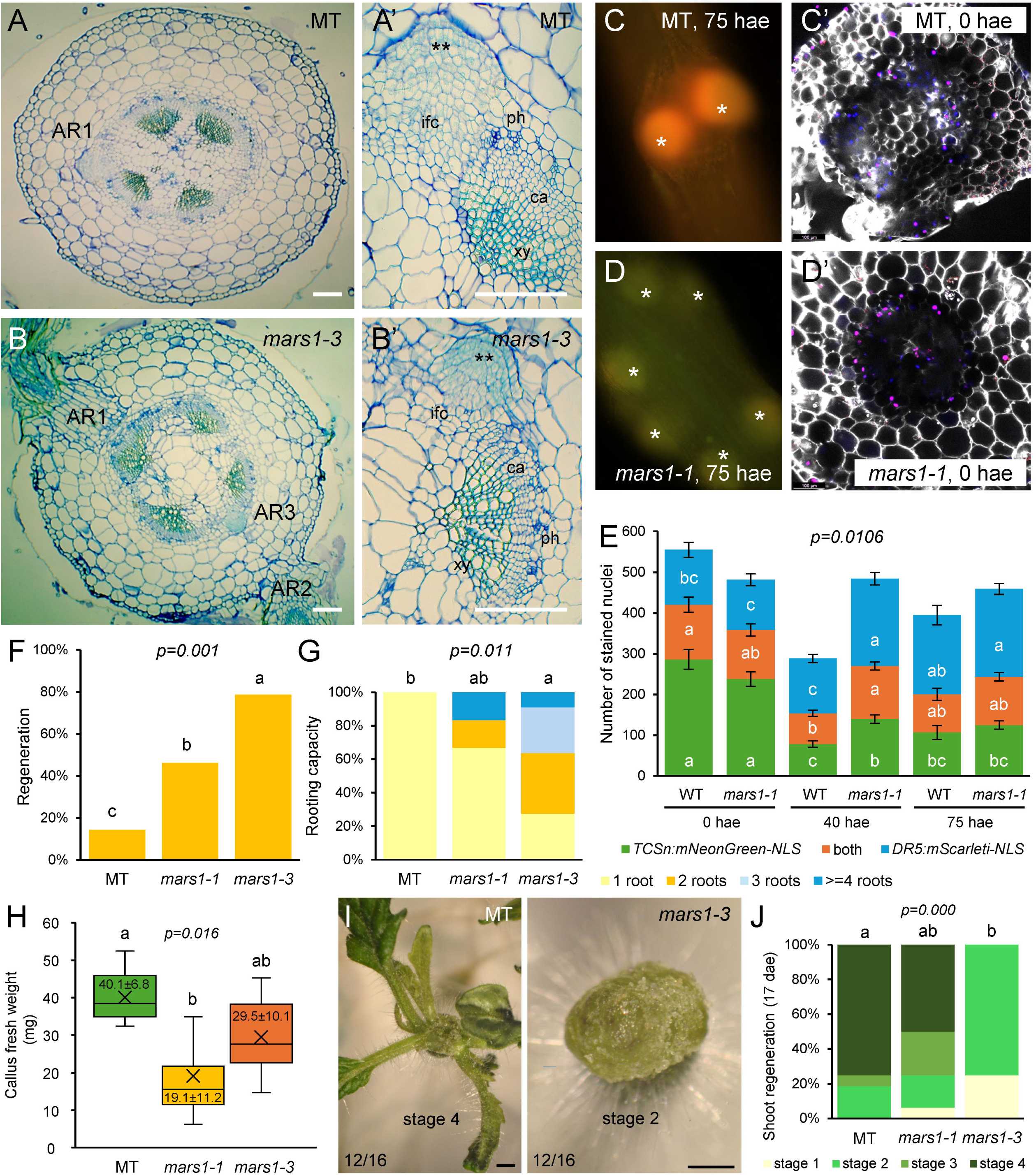
Morphological, anatomical and physiological changes in the basal region of hypocotyl explants in *mars1/rough* mutans during AR formation. (A-B) Representative cross sections of the basal region of the hypocotyl during AR formation in the MT (A, A’) and *mars1-3* mutant (B, B’). The AR primordia arise from the proliferation of the interfascicular cambium (ifc), on the side facing the phloem (ph) poles, and adjacent to the vascular bundle; c, cambium; xl, xylem. Double asterisks mark the quiescent center of the newest AR primordia. (C-D) Fluorescence microscopy and confocal images of MT (C, C’) and *mars1-1* (D, D’) hypocotyl sections expressing *DR5:mScarletI-NLS* (red) and *TCSn:mNeonGreen-NLS* (blue) markers (purple: overlap) at the different stages of AR development; C and D show the AR primordia protruding from the hypocotyl. (E) Quantification of the hormonal response in cross sections of the hypocotyl at different times during auxin-induced AR formation. (F) Regeneration percentage and (G) rooting capacity in cotyledon explants incubated with 0.15 µM NAA for 21 days. (H) Callus fresh weight in cotyledon explants incubated with 2.5 µM 6-BAP for 21 days. (I) Representative images of de novo organ formation in the apical region of hypocotyl explants at 17 days after excision (dae). (J) Shoot regeneration stages of hypocotyl explants at 17 dae as described in Larriba *et al*., (2021). Scale bars: 200 µm (A, B), and 1 mm (I). Different letters in E, F, G, H, J indicate significant differences between the studied samples (LSD; p-value<0.05; n>16).

Cotyledon explants of the *mars1/rough* mutants exhibit a heightened response to the exogenous addition of naphthaleneacetic acid (NAA), a synthetic auxin, as evidenced by their enhanced regeneration rate (**Fig. 4F**) and rooting capacity (**Fig. 4G**). Conversely, when cotyledon explants were incubated with the synthetic CK 6-benzylaminopurine (6-BAP), the *mars1/rough* mutants exhibited a diminished capacity for callus formation in comparison to their MT genetic background (**Fig. 4H**). These findings indicate that there has been a disruption in the endogenous signaling of the hormonal balance of auxins and CKs in the mutant tissues, as suggested by the results obtained with the fluorescent markers TCS and DR5 (see above). Subsequently, the extent of *de novo* shoot formation in the apical region of hypocotyl explants was evaluated in accordance with the methodology previously described (Larriba et al., 2021a). While a substantial proportion of MT hypocotyl explants exhibited the capacity to produce functional shoot apical meristems with fully developed leaves after three weeks (**Fig. 4I, J)**, the *mars1/rough* mutations significantly reduced the shoot regeneration capacity (**Fig. 4I, J)**. The *mars1-3* mutant exhibited a pronounced delay in *de novo* shoot regeneration, with most explants remaining in the stage of callus-like tissue formation (i.e., stage 2) for a longer time (**Fig. 4I, J**). These results agree with the enhanced auxin-to-CK response in the *mars1/rough* mutants, as stated above.

### Gene Expression Profile in the Base of the Hypocotyl Explants of *mars1/rough*

#### Mutants During the Early Stages of Adventitious Root Formation

Previous research has indicated that the gene regulatory networks involved in *de novo* organ formation in the basal region of the hypocotyl explants are differentially regulated in response to dynamic local hormone gradients established shortly after wounding (Larriba et al., 2021a; Larriba et al., 2021b; Larriba et al., 2022). Two independent RNA-Seq experiments with the *mars1/rough* mutants (**Supp. Fig. S4**) yielded expression data for 22,578 protein-coding genes (PCGs) and 1,206 long noncoding RNAs (lncRNAs) (**Table S2**, **Supp. Figs. S5**, and **S6**). These data represent 66.3% and 20.6%, respectively, of the tomato genes and lncRNAs in the ITAG4.0 genome annotation (Hosmani et al., 2019). Principal component analysis and k-means clustering of these datasets revealed substantial differences in transcript profiling over time and condition in both experiments, with less variation due to genotype (**Supp. Figs. S5A-D** and **S6A-D**). A total of 12,363 PCGs and 601 lncRNAs were identified as differentially expressed in the first experiment, regarding time (0 and 8 hae) and condition (shoot and hypocotyl explants) (**Table S3A**). Downregulated DEGs from the k-means analysis were found to be enriched in Gene Ontology (GO) terms related to photosynthesis (**Table S3B** and **Suppl. Fig. S5E**). In the second experiment, 9,073 genes that were deregulated in hypocotyl explants undergoing AR initiation at 20 and 48 hae were obtained (**Table S4A**). Several GO terms related to responses to oxidative stress were found to be significantly enriched among the upregulated DEGs identified from the k-means analysis in this experiment (**Table S4B** and **Suppl. Fig. S6E**). These results are consistent with the previously proposed working model for wound-induced AR formation in tomato hypocotyl explants (Larriba et al., 2021a; Larriba et al., 2021b; Larriba et al., 2022). Moreover, the present study builds upon this model by including the early response in AR formation in hypocotyl explants, both with and without a functional shoot apex.

Given the epigenetic function of SlLSD1, which is expected to broadly affect genome-wide gene expression, we employed a comparative approach to identify the genes that exhibited the most pronounced deregulation in the *mars1/rough* mutants. A total of 1,027 PCGs and 67 lncRNAs in the first experiment (**Table S3A** and **Suppl. Fig. S5F**), and 741 PCGs in the second experiment (**Table S4A** and **Supp. Fig. S6F**) were identified as being differentially expressed in the *mars1/rough* mutant hypocotyl explants compared to the MT background in the different time points and conditions (**Supp. Fig. S4**). Furthermore, 410 PCGs and 55 lncRNAs, and 446 PCGs were identified as deregulated in all mutant samples compared to the MT background in the first and second experiment, respectively (**Tables S3A** and **S3A**). A Venn diagram of these DEGs revealed that only 66 genes were deregulated in all contrasts, with most of them being upregulated in the *mars1/rough* mutants (**Table S5** and **Fig. 5A, B**). No significant enrichment of GO terms was observed among these DEGs. Our observations indicated that a significant proportion of these deregulated DEGs in the *mars1/rough* mutants were in genomic regions exhibiting low or residual expression levels in the MT background but flanked by genes exhibiting higher expression levels (**Fig. 5C** and **Supp. Fig. S7** and **S8**).

**Fig. 5.**
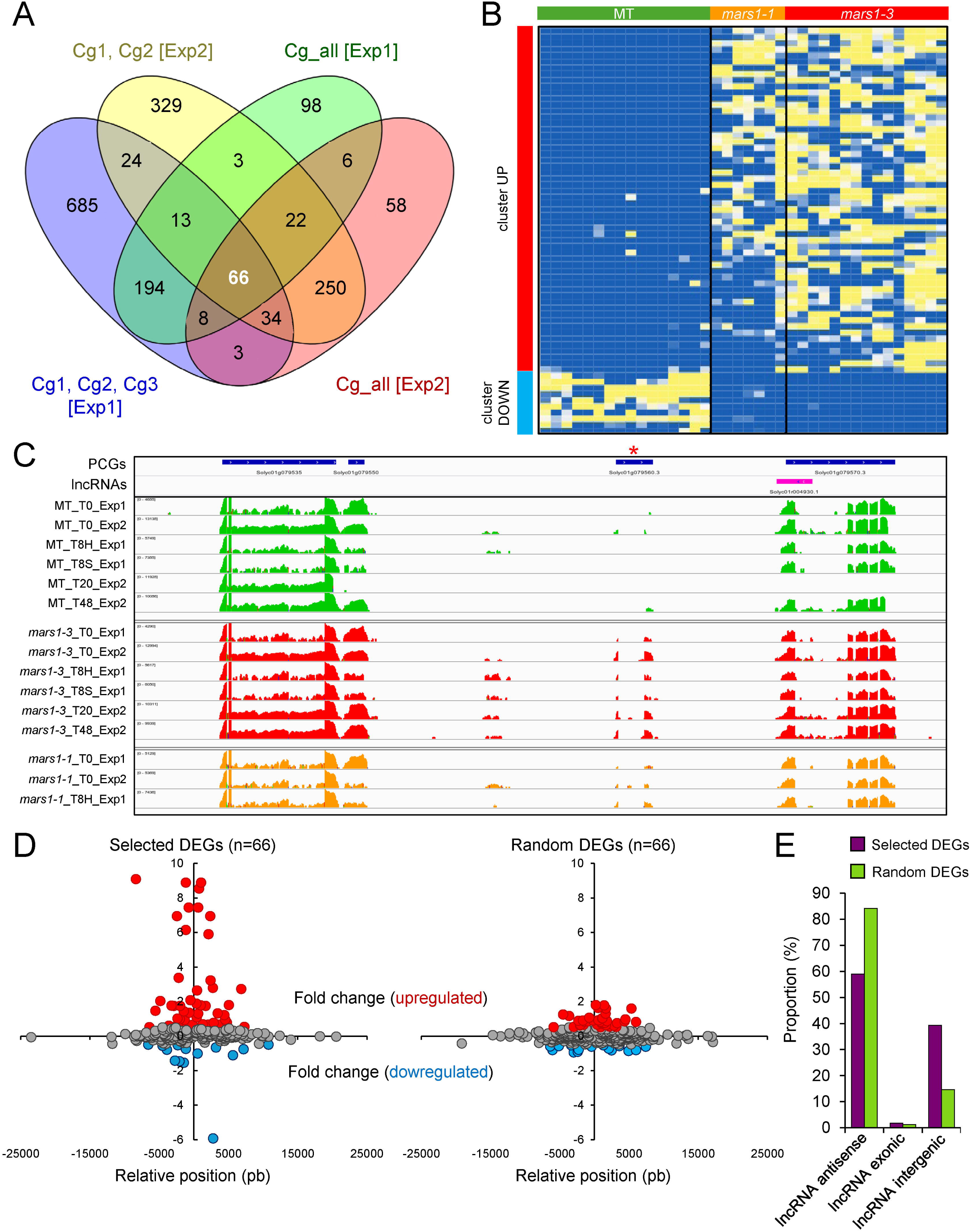
Identification of commonly deregulated genes in the *mars1/rough* mutants during AR formation. (A) Venn diagram of deregulated genes in the *mars1/rough* mutants compared to MT, as indicated in the main text and Suppl. Fig. S4A. Numbers indicate the DEGs found in each contrast (p-value<0.01). In white, selected DEGs that were studied further. (B) K-means clustering of selected DEGs. Upregulated and downregulated clusters are shown in red and blue, respectively (C) The colored peaks indicate the coverage of reads from multiple replicates at each time point, genotype, or condition. The MT samples are depicted in green, the *mars1-3* samples in red, and the *mars1-1* samples in orange. The annotated PCGs and lncRNAs are displayed at the top of each panel in blue and purple, respectively, with arrowheads indicating the direction of transcription. The red asterisks indicate genes within this region that were found to be significantly upregulated in the *mars1/rough* mutants compared to MT. (D) Deregulation of neighboring genes (50 Kb interval) of the commonly deregulated DEGs in the *mars1/rough* mutants compared to MT. (E) The transcripts annotated as lncRNAs in ITAG4.0 were then subjected to a hierarchical ranking based on their position in relation to protein-coding genes. The percentages were calculated based on the ratio of differentially expressed lncRNAs in the *mars1/rough* mutants compared to MT.

The presence of expressed PCGs and lncRNAs was determined in a 50 Kbp window centered on the selected 66 deregulated genes (see **Methods**). The degree of deregulation of the genes within these regions between the *mars1/rough* mutants and the MT background was visualized (**Fig. 5D**). These regions showed some enrichment for the presence of intergenic lncRNAs compared to their scattered distribution across the genome (**Supp. Fig. S7** and **Fig. 5E**). We extended this study to encompass all the other 557 DEGs identified in at least two contrasts (**Fig. 5A**). These results are consistent with the hypothesis that DEGs in *mars1/rough* mutants are mostly associated with regions containing intergenic lncRNAs, an association that may be functionally relevant for SlLSD1-mediated epigenetic regulation.

Our RNA-seq results indicate that SlLSD1 may play a pivotal role in the silencing of specific euchromatin regions, in a process that is likely facilitated by lncRNAs. To ascertain whether the *mars1/rough* mutations impact other non-coding regions of the genome, we have conducted *de novo* transcript annotation, which has enabled us to identify 27,985 expressed transcripts (see **Methods**; **Table S5**). The clustering of these transcripts during AR initiation according to their expression levels, from the lowest expression (first quartile, Q1) to the highest expression (fourth quartile, Q4), allowed us to identify that the most deregulated genes in the *mars1/rough* mutants were included among genes in Q1 and, to a lesser extent, in Q2 (**Fig. 6A, B**). Upon closer examination of the deregulated transcripts in Q1 in the mutants, it became evident that these transcripts exhibited a broad distribution along the entire chromosome, extending beyond the euchromatic arms to encompass the centromeric and pericentromeric chromatin (**Fig. 6C**). Conversely, deregulated genes in the mutants assigned to Q2 and Q3 were predominantly located within euchromatin regions of the chromosome arms (**Fig. 6C**). The deregulated transcripts in the *mars1/rough* mutants have been annotated according to their functional characteristics. Of interest is the observation that a significant proportion of the deregulated transcripts in Q1 are in intergenic regions where no transcripts have been previously assigned (**Fig. 6D**).

**Fig. 6.**
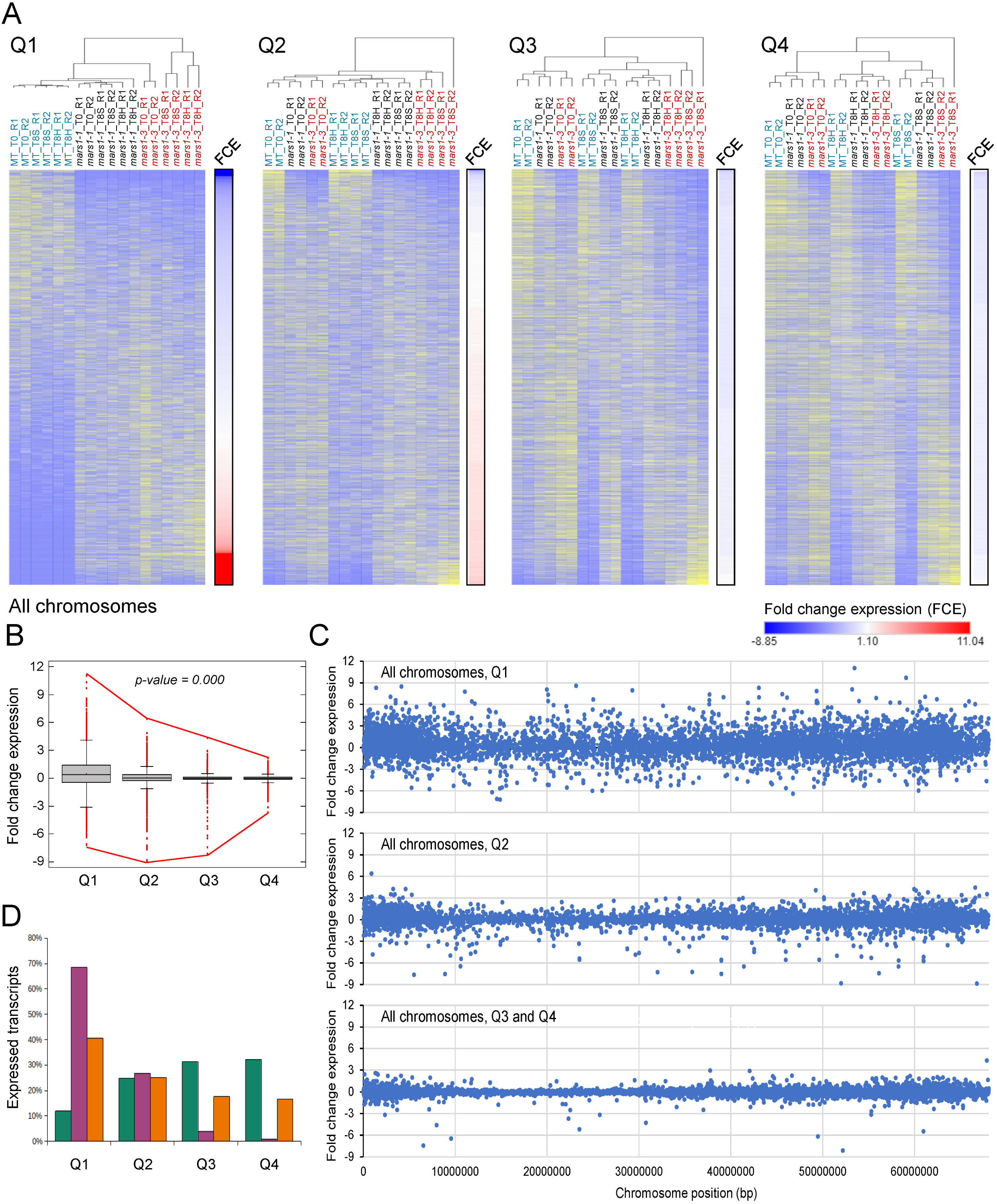
Transcriptome landscape of *mars1/rough* mutants during AR initiation. (A) Clustering of expressed transcripts at the base of the hypocotyl during AR initiation according to their expression levels, from the lowest expression (first quartile, Q1) to the highest expression (fourth quartile, Q4). The clustering of rows was conducted according to the fold change in expression between the mutants (*mars1-1* and *mars1-3*) and the MT datasets. The clustering of columns was conducted using the Euclidean distance between the different libraries. The relative expression levels in each row are presented on a scale of 0 (blue) toL1 (yellow). The fold change in expression ranges from −8.85 (blue) to 11.04 (red). (B) Box plot of fold change expression of the expressed transcripts in the studied quartiles. The red lines connect the maximum and the minimum values in each quartile. The p-values are indicated, with those in italics indicating statistical significance (T-test; p-value<0.05; n=6,697). (C) Fold change expression of the expressed transcripts along the chromosome position, in bp. (D) Functional classification of transcriptional regions, according to whether they contain at least one PCG or lncRNA as it has been annotated in ITAG4.0 (green), intergenic transcriptional regions that do not contain a PCG or lncRNA (purple), or with transcriptional units not included in any of the above (orange).

#### SlLSD1 is Involved in the Maintenance of Monomethylated Histone H3 Lysine 4

In metazoans, LSD1/KDM1A demethylates the mono- or di-methylated histones H3 lysine 4 (H3K4; Shi et al., 2004) and H3K9 (Metzger et al., 2005). To ascertain which specific methylation modifications were significantly altered by the *mars1/rough* mutations, we conducted immunoblot analysis with several antibodies to analyze the abundance of modified H3 proteins (see **Methods**). The levels of H3K4me1 proteins were found to be significantly higher in the *mars1-3* mutants compared to those of the MT background (**Fig. 7A, B**). The levels of histone H3K4me2 proteins also appeared to be higher in the *mars1-3* mutants than in the MT plants (**Fig. 7A, B**). In contrast, no differences in the levels of H3K4me3 proteins were observed between MT and *mars1-3* mutants (**Fig. 7A, B**). The levels of two additional histone H3 marks associated with transcriptional activation, H3K36me3 and acetylated H3K9 (H3K9Ac; Berr et al., 2011), were quantified, with no statistically significant differences observed between MT and *mars1-3* mutants (**Fig. 7C, D**). These findings provide evidence that supports the hypothesis that the tomato SlLSD1 protein regulates the demethylation of H3K4me1.

**Fig. 7.**
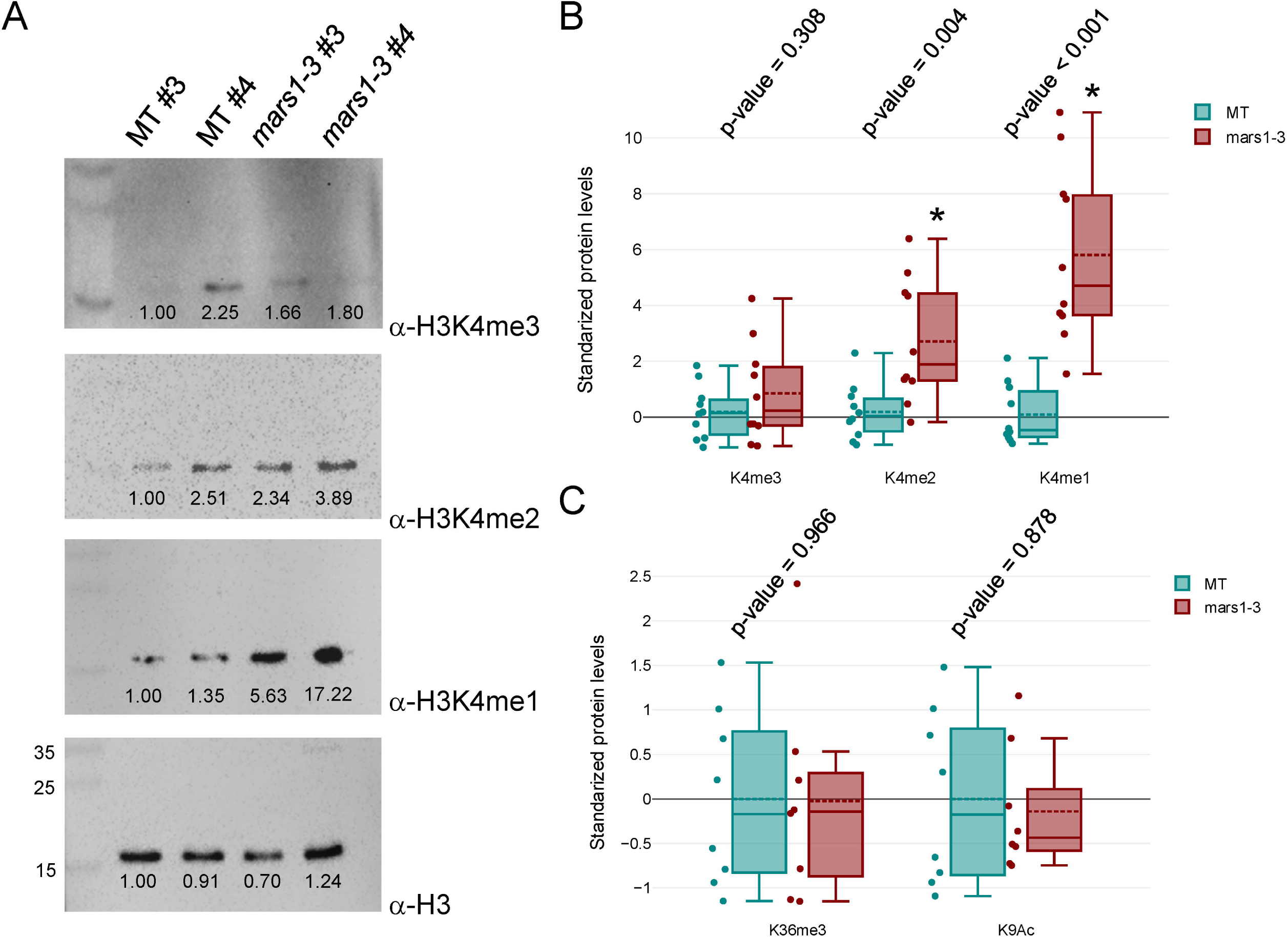
Several histone marks were found altered in the *mars1/rough* mutants. (A) Representative Western blotting images of some H3 marks in proteins extracted from two independent MT and *mars1-3* mutant plants. Numbers on the membrane indicate relative quantification regarding MT #3. Numbers in the left corner indicate the size, in Kb, of the protein marker. (B, C) The alterations of H3K4me3, H3K4me2, H3K4me1, H3K36me3 and H3K9Ac were normalized with internal H3 protein signals. The experiment was repeated at least four times. The histogram illustrates the levels of modified H3 proteins in MT and *mars1-3* mutants. Asterisks indicate highly significant differences as regards the MT background (T-test; p-value <0.01).

#### Functional Validation of the Gene Supercluster Responsible for the *mars1/rough* Phenotype

A search of the deregulated regions in the *mars1/rough* mutants for genes whose orthologs in other species are involved in *de novo* organ formation during regeneration (Larriba et al., 2022) yielded no results. Among the regions showing transcript upregulation in the *mars1/rough* mutants, a ∼100 Kbp region on chromosome 4 (SL4.0ch04: 63,531,701 – 63,631,100) was identified as a potential candidate region for further investigation (**Suppl. Fig. S7** and **Fig. 8A**). The antisense lncRNA *Solyc04r023410* located in this region was found to be upregulated in the *mars1/rough* mutants, together with several other RNAs spanning the coding sequences of three contiguous genes: *Solyc04g081650*, *Solyc04g081660* and *Solyc04g081670* (**Fig. 8B**). Our directional RNA-seq allowed us to reconstruct seven different transcripts including these three genes (**Fig. 8A**). *Solyc04g081660* and *Solyc04g081650* have been annotated to putatively encode proteins with partial homology to the N- and C-terminal domains of mitotic B-type cyclins, respectively. Both genes were found to be expressed at low levels in different tissues and over developmental stages in the ‘M82’ background, except in generative and sperm cells during pollen development (**Suppl. Fig. S9**). We confirmed by RT-PCR that several transcripts on this region were upregulated in both young shoot explants and fruit skin in the *mars1-1* and *mars1-3* mutants, with no detectable expression in their corresponding MT background (**Fig. 8C** and **Table S7**). We cloned and sequenced five of the transcripts encompassing the *Solyc04g081650-Solyc04g081670* genomic region, which showed differential expression in shoot explants and fruit skin tissue (**Fig. 8D**). Due to the absence of expression of these transcripts in the MT background in our RNA-seq (**Fig. 8B**), we hypothesize that the higher expression of some of these transcripts in the *mars1/rough* mutants may contribute to their enhanced AR formation in the hypocotyl after wounding and uncontrolled cell proliferation in the fruit skin. To check the genomic structure including these two candidate genes in the MT and *mars1/rough* mutants, a 3.6 Kbp region was sequenced and compared with the MT reference genome (Shirasawa and Ariizumi, 2024). Only four mismatches were found between the ‘Heinz 1706’ reference genome and MT, *mars1-1* and *mars1-3* mutants (**Supl. Fig. S10A**). We found many SNPs and InDels when comparing this region with the homologous region encompassing *Solyc04g082430*, located 477 Kbp away on the same chromosome (**Supl. Fig. S10A**). Interestingly, these two genomic regions were highly conserved in close tomato relatives such as *S. pimipinellifollium* and *S. pennellii* (**Supl. Fig. S10B**), suggesting a common origin of both genomic sequences in the common ancestor of these three species, probably due to an ancient genomic duplication. In silico translation of *Solyc04g081670*-*Solyc04g081650* transcripts identified two peptides that partially encode an incomplete B-type cyclin, closely related to the G2/mitotic-specific cyclin B2;4 encoded by *Solyc04g082430* (**Fig. 8E**). The peptide encoded by *Solyc04g081650* encompassed the canonical C-terminal domain of the cyclin proteins (PF02984), with characteristic alpha-helixes fully overlapping with the C-terminal domain of cyclin B2;4 (**Fig. 8F**). The peptide encoded by *Solyc04g081660* showed some similarity to the N-terminal region of the B-type cyclins, but lacked most of the internal amino acids present in the canonical N-terminal domain sequence (PF00134) (**Fig. 8E**). B-type cyclins are regulated by APC/C-mediated ubiquitination through their DEAD-box (Davey and Morgan, 2016), and we found a sequence matching the consensus DEAD-box in the N-terminal peptide that is expressed in the *mars1/rough* mutants (**Fig. 8E**).

**Fig. 8.**
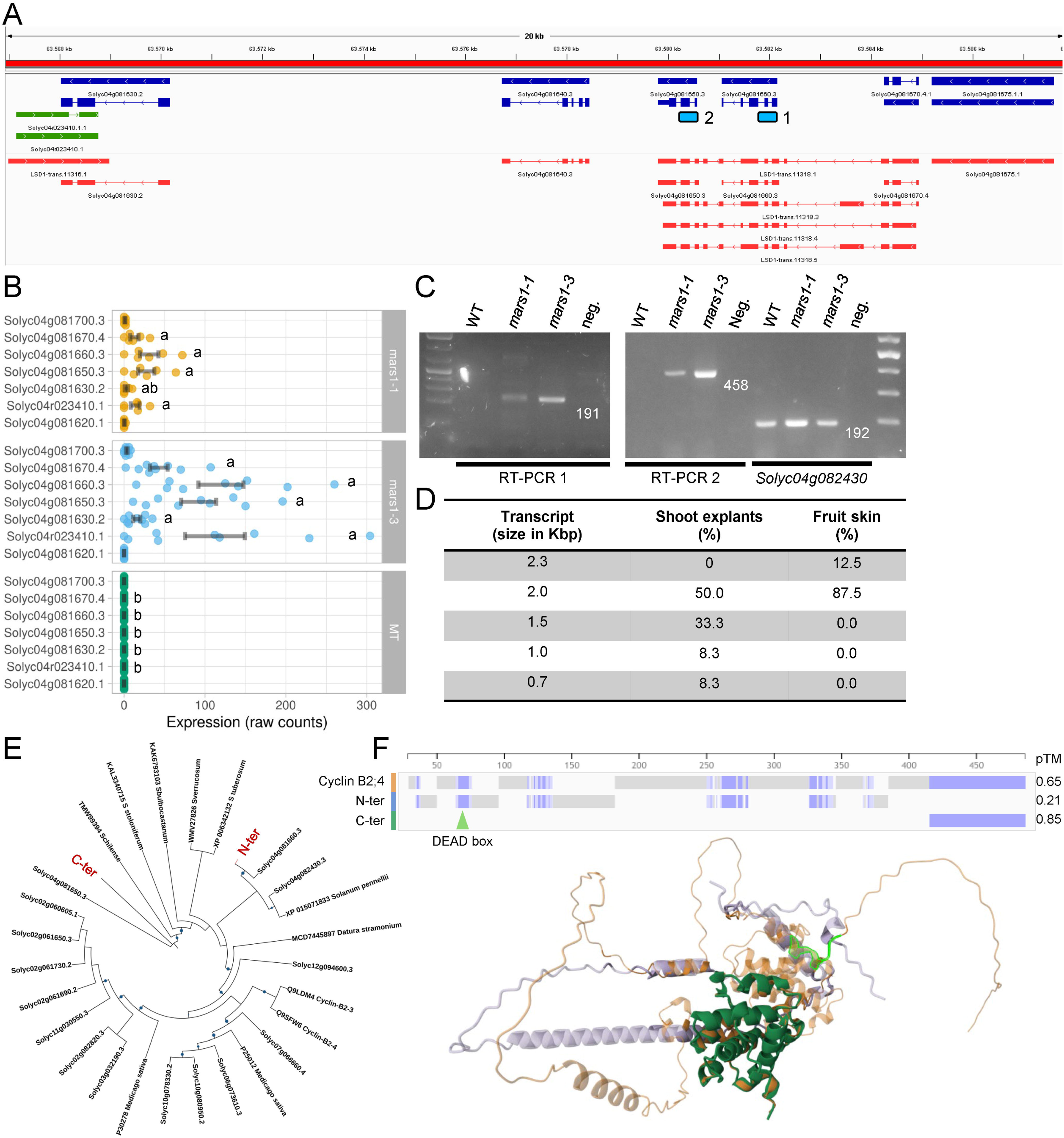
Functional validation of the gene supercluster responsible for the *mars1/rough* phenotype. (A) A 20 Kbp region corresponding to the gene supercluster that is highly deregulated in *mars1/rough* mutants. The genes and lncRNAs annotated in ITAG4.0 are shown in dark blue and green, respectively, and the transcripts predicted from our RNA-Seq experiments are shown in red. Light blue rectangles indicate products (1, 2) obtained from RT-PCR. (B) Expression data (in raw counts) of the six genes and one lncRNA included in this region obtained from our RNA-seq (experiment 1). Letters indicate highly significant differences between genotypes for a given gene (T-test; p-value <0.01). (C) RT-PCR of hypocotyl samples at time 0 (T0). RT-PCR 1 and 2 correspond to partial amplification of the annotated genes *Solyc04g081660* and *Solyc04g081650*, respectively. RT-PCR of *Solyc04g082430* was used as a control for cDNA quality. Water was used as a negative control (neg.). The numbers indicate the expected size of the amplified bands from the cDNA. (D) Novel transcripts expressed in *mars1/rough* mutants. Percentage was estimated based on the number of clones identified for each transcript. (E) Phylogenetic tree of the closest relatives to cyclin B peptides encoded by *Solyc04g081660* (N-ter), and *Solyc04g081650* (C-ter). The protein sequences were aligned using T-Coffee, and a phylogenetic tree was built with IQ-TREE2, employing the JTT-G4 model and iToL representation. (F) Alignment of 3D structures of peptides encoded by *Solyc04g082430* (cyclin B2;4, orange)*, Solyc04g081660* (N-ter, purple), and *Solyc04g081650* (C-ter, green). The putative DEAD-box domain is indicated in light green. Predicted template modeling (pTM) score of the 3D structure shown is indicated.

To corroborate the hypothesis that the *Solyc04g081670-Solyc04g081650* locus is involved in the *mars1/rough* phenotype, we employed a virus-induced gene silencing (VIGS) strategy (Fu et al., 2005) to suppress the expression of this gene in the *mars1/rough* mutants (see **Methods**). Agro-injection of immature fruits (8-10 DPA) of the *mars1/rough* mutants was performed, and it was determined that the region surrounding the injection does not exhibit the roughness that characterizes the epidermis of mature fruits of the *mars1/rough* mutants (**Fig. 9A, B**), whereas the VIGS of the endogenous *PHYTOENE DESATURASE* (*PDS*) gene reduced pigmentation of mature *mars1/rough* fruits without modifying their roughness (**Fig. 9A, B**). This can be attributed to the absence of ectopic proliferation in the targeted region following VIGS induction. RT-PCR analysis has been employed to confirm the expression of VIGS and the downregulation of the target RNAs in these fruits. Our results demonstrate that several transcripts are upregulated in the *mars1/rough* mutants in a tissue-specific manner, some of which may encode a truncated mitotic cyclin B protein that may interfere with endogenous cyclin B protein function (**Fig. 9C**). The accumulation of a putative non-degradable cyclin may induce ectopic cell divisions, as it has been reported in other plant species (Weingartner et al., 2004).

**Fig. 9.**
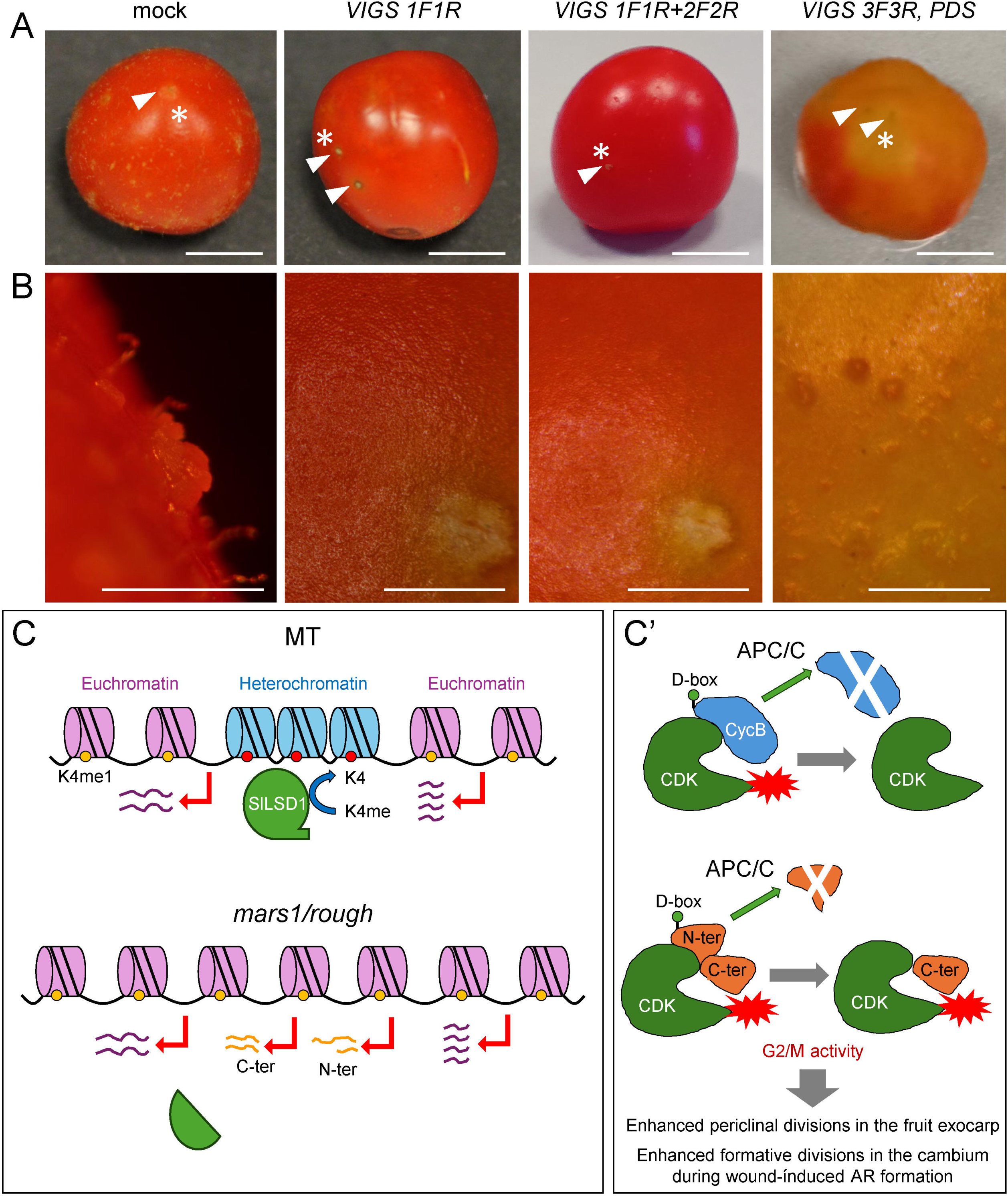
Downregulation of *Solyc04g081650* and *Solyc04g081660* genes using a virus-induced gene silencing (VIGS) approach. (A) Representative images of mature fruits of *mars1-3* mutants 14 days after being injected with *Agrobacteriun tumefaciens* strains containing pTRV1 or pTRV2-GG, either empty (mock) or with *Solyc04g081650* (*VIGS 1F1R*), *Solyc04g081660* (*VIGS 2F2R*) or *Solyc03g123760* (*VIGS 3F3R*, *PDS*) probes, respectively. White arrowheads in A indicate the injection points. (B) Details of the fruit surface displaying tissue outgrowth in these fruits *mars1/rough* mutants. Insets correspond to the regions indicated in A with white asterisks. (C, C’) Proposed model for SlLSD1 regulation of chromatin and cell cycle control. Scale bars: 10 mm (A), and 1 mm (B).

## Discussion

Two independent mutant screenings revealed that the altered function of *Solyc10g047350* results in the manifestation of mild developmental phenotypes. This gene encodes a putative demethylase of histone marks H3K4me1 and H3K9me2, homologous to LSD1/KDM1A proteins, henceforth designated *SlLSD1*. The altered function of *SlLSD1* produced a pleiotropic phenotype noticeable throughout the tomato life cycle. The mutations in this gene affected different traits, including early seedling growth, AR formation, inflorescence architecture, fruit exocarp development, and embryonic development. Based on the AR and fruit pericarp phenotypes, we have dubbed these mutants as *mars1*/*rough*. Moreover, it has been observed that some of the traits under study exhibit an enhancement of the mutant phenotypes in successive generations, which might be related to the epigenetic function of SlLSD1. This observation is consistent with previous findings in the Arabidopsis *fasciated* mutants with disrupted nucleosome assembly, which display transgenerational aggravation of their phenotypes associated with several alterations in DNA methylation at transposable elements (TEs) (Mozgova et al., 2018). Further studies on the *mars1/rough* mutants in successive generations are necessary to elucidate the mechanism behind the progressive worsening of their mutant phenotype.

The size of tomato fruit is determined by the relative contribution of cell division and cell expansion, which exhibit specificity at both the tissue and temporal levels (Renaudin et al., 2017). The pericarp tissues of the fruit are derived from the carpel walls and are composed of three layers of exocarp (E1 to E3), the central layers of mesocarp, and two inner layers of endocarp (I1 and I2). During early fruit development, the exocarp is the primary site of cell division, while cell expansion takes place mainly in the mesocarp, in close association with the progression of endoreduplication (Tourdot et al., 2024). Auxin and gibberellins play an important role in the coordination of cell division and expansion during the early stages of fruit development (Pattison and Catalá, 2012; Renaudin et al., 2023), and it is well known that endogenous CKs regulate pericarp cell division (Gan et al., 2022). The majority of the divisions are anticlinal in the epidermal layer, while periclinal divisions are observed in the underlying sub-epidermal cells, giving rise to the formation of additional mesocarp layers (Renaudin et al., 2017). In the fruit epidermis of *mars1/rough* mutants, abnormal cell division patterns, absent in the ovary at anthesis, occur in the exocarp as soon as 5 DPA. The fruit epidermis, normally consisting of a single layer of cells with anticlinal divisions, undergoes periclinal divisions in *mars1/rough* mutants, which are normally largely limited to the E2 and E3 sub-epidermal cell layers. As a result, supernumerary cell layers appear in the fruit epidermis, eventually leading to the appearance of protuberances on the fruit surface. Later, during the cell expansion and following ripening stages, the uncontrolled cell division in the epidermis leads to the disruption of fruit skin in the protruding areas.

The observed fruit phenotype in the *mars1/rough* mutants is consistent with the hypothesis that a temporal dysregulation of cell division and/or cell division plane determination occurs in some cells of the fruit epidermis. B-type cyclins are among the primary regulators of the G2/M transition, which is critical for the regulation of cell division and, consequently, fruit development. The results of this study suggest that several peptides with incomplete cyclin B features, potentially encoded by the *Solyc04g081660*-*Solyc04g081650* locus, were specifically upregulated in the *mars1/rough* mutants. These peptides may play a role in the development of the rough fruit phenotype. The constitutive overexpression of a non-degradable cyclin B1 protein in transgenic tobacco seedlings resulted in strong developmental abnormalities due to impaired cell cytokinesis, which led to the formation of polynucleated cells (Weingartner et al., 2004). Conversely, the ectopic overexpression of APC10, one of the key subunits of the anaphase-promoting complex/cyclosome (APC/C) that functions to degrade mitotic cyclins at the G2/M transition and exit from mitosis (Vodermaier, 2004), was found to enhance the rates of cell division during the early stages of leaf development, without affecting plant morphology (Eloy et al., 2011). However, the cellular phenotype of *mars1/rough* cell clusters in fruits, resembles the ectopic constitutive expression of *CYCD3;1* in Arabidopsis seedlings, which resulted in a shortened G1 phase and inhibited cell cycle exit, which in turn led to smaller cells and ectopic divisions in many different tissues through regulation of the RETINOBLASTOMA RELATED-EF2 canonical pathway (Dewitte et al., 2003). We hypothesize that the novel B-type cyclin peptides that are upregulated in the *mars1/rough* mutants increase the mitotic rate of some epidermal cells in a cell-autonomous manner, independently of their regulation by the APC/C. This finding aligns with the short-range activation of irregular divisions observed in the epidermis of *mars1/rough* fruits.

In tomato fruits, the cell expansion phase, after 10−12 DPA, is associated with a considerable increase in the thickness of the fruit cuticle, which is formed by epidermal cells (Reynoud et al., 2023). One of the upregulated genes in the *mars1/rough* mutants was *CUTIN DEFICIENT 2* (*CD2*), *Solyc06g035940*, which encodes a member of the class IV homeodomain-Leu zipper (HD-ZIP IV) family (Nadakuduti et al., 2012). Mutations in *CD2* have been found to cause a significant reduction in the biosynthesis of cutin in tomato, the main cuticle polymer, and altered wax deposition (Isaacson et al., 2009; Nadakuduti et al., 2012). Accordingly, while wax load and composition were not altered in *mars1/rough* mutants, the cutin load was considerably increased, suggesting the coordinated up-regulation of genes involved in cuticle polymer formation, as observed in other tomato cutin-enriched mutants (Petit et al., 2021). Moreover, the composition of *mars1/rough* cuticle lipid polyesters exhibited the signature of suberin, a cuticle polyester covering underground plant organs and seeds. As suberin is also synthesized in aerial organs for wound-healing, as shown in tomato fruit cuticle mutants (Lashbrooke et al., 2016), this suggests that suberin is synthesized in response to the disruption of the fruit epidermis, giving the *mars1/rough* fruit its russet-colored aspect. Consequently, the complex and pleiotropic phenotype observed in the *mars1/rough* mutants might be a consequence of altered expression of several genes which are normally silenced through SlLSD1-mediated H3K4me1 demethylation.

In Arabidopsis, the primary methyltransferase responsible for H3K36me2/3-mediated transcriptional activation is SET DOMAIN GROUP 8 (SDG8). Its deficiency results in a range of pleiotropic developmental abnormalities, including a reduction in wound-induced AR formation in leaf explants (Zhang et al., 2019). In their working model, SDG8 function is required to adequately transduce the wound signal to upregulate *ANTHRANYLATE SYNTHASE* α*1* (*ASA1*), a tryptophan biosynthesis gene in the auxin production pathway, which is required to trigger formative cell divisions in the cambium cells (Zhang et al., 2019). Moreover, the formation of callus in response to hormones is contingent upon the stimulatory effect of auxin on the upregulation of SDG8, which in turn determines the status of H3K36me3 on callus-related genes including *CYCLIN B1;1* (Ma et al., 2022). Furthermore, ARABIDOPSIS TRITHORAX-RELATED 2 (ATXR2) is a histone methyltransferase that facilitates the accumulation of H3K36me3 at the promoters of *LATERAL ORGAN LIMITED DOMAIN* (*LBD*) genes, which are activated during cellular dedifferentiation in response to elevated levels of endogenous auxin during hormone-triggered regeneration. ATXR2 is recruited to *LBD* promoters through its interaction with the AUXIN RESPONSE FACTOR 7 (ARF7) and ARF19 transcription factors (Lee et al., 2017). These results indicate that auxins act both upstream and downstream of epigenetic regulators such as SDG8 and ATRX2, which may complicate the interpretation of the pleiotropic phenotypes observed in these epigenetic mutants. In the tomato *mars1/rough* mutants, which showed altered response to the hormonal balance of auxins and CKs, no alterations were observed in global levels of H3K36me3 nor any deregulation in the orthologs of genes directly involved in regeneration in other species. Hence the enhanced wound-induced AR formation in hypocotyl explants and ectopic callus formation phenotype in fruits may rely on a distinct regulatory mechanism requiring SlLSD1.

According to the working model depicted in **Fig. 9C**, SlLSD1predominantly functions as a histone demethylase, specifically targeting H3K4me1 in various regions of constitutive heterochromatin situated within euchromatin-rich chromosome arms. These regions are distinguished by the presence of intergenic lncRNAs and are abundant in TEs and pseudogenes. In SlLSD1-deficient *mars1/rough* mutants, an increase in gene expression is observed in these regions, leading to the accumulation of novel transcripts, some of which have coding potential. One region with significant deregulation in *mars1/rough* mutants contains a pseudogene annotated as two contiguous genes *Solyc04g081660* and *Solyc04g081650*. This pseudogene encodes at least two peptides that are homologous to the N- and C-terminal regions of *Solyc04g082430*, respectively. *Solyc04g082430* encodes the G2/M-specific SlCycB2;4, which is expressed during gametogenesis. In the reprogramming system for callus formation from protoplasts obtained from the leaf mesophyll in Arabidopsis (Sakamoto et al., 2022), it has been observed that the biosynthesis of endogenous auxin induces the transcriptional reactivation of cell cycle genes from the G2/M phase, including *B-TYPE CYCLIN-DEPENDENT KINASES* and *cyclin B* genes, among others (Sakamoto et al., 2022). Indeed, mutations in *CYCD3;1* and its homologs *CYCD3;2* and *CYCD3;3* cause defects in callus formation at wound sites in Arabidopsis hypocotyl explants (Ikeuchi et al., 2017). Given the structural similarity between the two peptides that are constitutively expressed in *mars1/rough* mutants and SlCycB2;4, it is hypothesized that they interact with cyclin-dependent kinases (CDKs). In tomato, two genes encoding CDKB1 and CDKB2 proteins are predominantly expressed in the pericarp during the early developmental stages, when cell division takes place (Czerednik et al., 2012). CDKA1 is expressed in the pericarp throughout development but is strongly upregulated in the outer pericarp cell layers at the end of the growth period, when *CDKB* gene expression has ceased. Overexpression of the *CDKB* genes at later stages of development and the downregulation of *CDKA1* result in a very similar fruit phenotype, showing a reduction of endoreduplication and in their cuticle thickness (Czerednik et al., 2012). The cyclin B-CDK interaction has been demonstrated to positively regulate CDK kinase activity, thereby enhancing mitotic progression in these tissues. However, the DEAD-box, necessary for cyclin B regulation via APC/C-mediated ubiquitination, is only present in the N-terminal peptide. Consequently, the prolonged binding of the C-terminal peptide to CDKA1 in the *mars1/rough* mutants could maintain their kinase activity for a longer period and induce the increased proliferation observed in the fruit exocarp and cambial hypocotyl during wound-induced AR formation. This proliferation is likely correlated with high endogenous auxin levels. A comparable mechanism may function in other *mars1/rough* organs, such as leaves, but other established mitotic regulators, such as INHIBITOR OF CDK/KIP-RELATED PROTEIN (ICK/KRP) (Nafati et al., 2010) or WEE1 protein kinase (Gonzalez et al., 2007), might counteract cyclin B/CDK activity in these organs.

Despite the upregulation of a putative mitotic cyclin in *mars1/rough* mutants, transcriptomic analyses of tomato hypocotyl explants during wound-induced AR formation indicate that *mars1/rough* mutations alter the regulation of several dozen silenced genomic regions. Our findings indicate that SlLSD1 appears to act locally in the silencing of specific regions located within euchromatin in the chromosome arms, and globally in the silencing of regions of pericentromeric heterochromatin. In both cases, the presence of lncRNAs in these regions appears to be associated with SlLSD1-mediated gene regulation. In Arabidopsis, DNA methylation is correlated with specific histone marks (Zhang et al., 2009; Roudier et al., 2011). Global DNA methylation during the hormone-induced leaf-to-callus transition was investigated in this species (Shim et al., 2022). Key cell division genes showed decreased DNA methylation in callus tissue, which correlated with their increased gene expression in this tissue. This suggests that dynamic changes in DNA methylation activate cell division genes during the leaf-to-callus transition, likely in response to the hormonal balance between auxin and CKs. The alterations observed in gene-body regions were relatively minor and consistent across all DNA methylation contexts throughout the leaf-to-callus transition. In contrast, dynamic alterations in DNA methylation were observed primarily in regions containing transposon TEs, which is consistent with the main role of DNA methylation in maintaining genome stability. The loss-of-function mutations of two partially redundant members of the LYSINE-SPECIFIC DEMETHYLASE 1-LIKE (LDL) family in Arabidopsis, *LDL1* and *LDL2*, result in a reduction in RNA-directed DNA methylation (Greenberg et al., 2013). A number of studies have reported the genome-wide occupancy profile of LDL1/2, which are known to interact with the histone deacetylase HDA6 to synergistically regulate gene expression by removing H3Ac and H3K4me1/2 in their target genes (Hung et al., 2018; Hung et al., 2020). It is noteworthy that the findings indicated the potential involvement of HDA6-LDL1/2 in the regulation of a specific subset of lncRNAs, which exhibited low or undetectable expression in wild-type Arabidopsis seedlings. Moreover, the expression levels of the neighboring genes were found to be positively correlated with the expression of these lncRNAs (Hung et al., 2020). It can thus be postulated that HDA6-LDL1/2 may play a pivotal role in plant development by repressing some lncRNAs that may be induced under specific environmental conditions. A recent report also shed some light in the synergistic regulation of HDA6-LDL1/2 in TE de-repression likely through DNA hypomethylation (Hsieh et al., 2024).

Additionally, the deregulation of additional pericentromeric regions associated with TEs was observed in *mars1/rough* mutants. In Arabidopsis, several H3K9 methylase genes, including *SU(VAR)3–9-RELATED 5* (*SUVR5*) and *KRYPTONITE (KYP)/SUVH4*, *SUVH5* and *SUVH6*, are known to be involved in H3K9me2 deposition in TEs in a DNA methylation-independent and DNA methylation-dependent manner, respectively (Caro et al., 2012; Du et al., 2014). The loss of H3K9me2 in TEs, as observed in *suvh456* triple mutants, induces a gain of H3K4me1, which in turn leads to their transcriptional activation (Inagaki et al., 2017). Among the four LSD1Llike genes in Arabidopsis, *LSD1-LIKE 2* (*LDL2*) has been proposed to function downstream of H3K9me2, most likely by decreasing H3K4me1 levels (Inagaki et al., 2017). Furthermore, SUVR5 has been demonstrated to interact physically and genetically with LDL1, another H3K4 demethylase that is partially redundant with LDL2 (Caro et al., 2012). The loss of KYP histone methyltransferase in tomato revealed that the levels of H3K9me2 in *kyp* mutants diminished in discrete pericentromeric regions, which in turn showed a sharp deposition of H3K9 acetylation and the emergence of novel topologically associating domains (TAD)-like structures that could contribute to chromatin 3D conformation and changes in KYP-controlled gene expression (An et al., 2024). Therefore, we cannot rule out the possibility that SlLSD1 is involved in the epigenetic regulation of intercalated heterochromatin through a pathway like that proposed for LDL2 in Arabidopsis (Inagaki et al., 2017).

Our results suggest that SlLSD1, which plays a similar role to that of its Arabidopsis homologs, LDL1 and LDL2, is responsible for maintaining genomic integrity by repressing specific target genes interspersed within euchromatin. The use of tagged SlLSD1 proteins in conjunction with ChIP-seq experiments utilizing the H3K4 marks that are deregulated in the *mars1/rough* mutants will facilitate the identification of the specific genomic regions that are regulated by SlLSD1 and its interacting protein partners.

## Methods

### Plant Materials and Growth Conditions

The seeds of *Solanum lycopersicum* L. cv. ‘Micro-Tom’ and the P19E6 line, obtained from a highly mutagenized EMS mutant collection (Just et al., 2013), were utilized in all experiments. A total of thirty M_2_/M_3_ mutant families were selected through the screening of our phenotypic mutant database (Just et al., 2013) for the presence of fruit cracking mutants. The reporter line *DR5:mScarletI-NLS*;*TCSn:mNeonGreen-NLS*. has been previously described (Omary et al., 2022). The seeds were surface-sterilized and germinated *in vitro* in accordance with the methodology indicated by Alaguero-Cordovilla et al. (2020). Following this, the one-month-old seedlings were transplanted and cultivated in the greenhouse under standard conditions, as described by Rothan et al. (2016).

### Morphological Analysis of *mars1/rough* Phenotypes

The early seedling phenotype of *mars1/rough* mutants was studied as indicated by Alaguero-Cordovilla et al. (2020). The formation of ARs was induced by removing the entire root system 2–3 mm above the hypocotyl-root junction of *in vitro* grown tomato seedlings at the 100–101 growth stages, and the shoot explants were subsequently transferred to glass jars containing standard growth medium (Alaguero-Cordovilla et al., 2021). Each jar contained six explants of the same genotype and/or treatment. All experiments were conducted in triplicate. Additionally, hypocotyl explants were obtained by sectioning the shoot explants just below the cotyledons. The emergence of ARs from the basal region of the hypocotyl explants was monitored and recorded at regular intervals. The emergence of ARs was estimated based on the day before the first AR was observed. The regeneration of shoot tissue in hypocotyl explants was characterized according to the methodology described by Larriba et al. (2021a).

To assess the impact of exogenous hormones on the process of tissue regeneration, cotyledon explants were harvested from 7-day-old seedlings and divided into two halves. Subsequently, the explants were incubated in 90-mm diameter Petri dishes containing Murashige and Skoog (MS) basal salt medium with 2% sucrose, 0.8% agar with or without the addition of 0.15 µM NAA or 2.5 µM 6-BAP. After 21 days of incubation, the number of ARs and the fresh weight of callus were quantified. The experiments were conducted in triplicate with a minimum of six explants used in each trial.

### Mapping-by-Sequencing and Recombinant Analysis

Mapping-by-sequencing and recombination analysis were performed as previously described (Petit et al., 2016; Garcia et al., 2016). A mapping population (BC_1_F_2_) was created by crossing the P19E6 mutant with a MT parental line and by selfing a single BC_1_F_1_ hybrid. Mutant bulk was then constituted by pooling 44 plants displaying a rough fruit phenotype and DNA extracted was used for preparation of library that was sequenced using a HiSeq 2500 sequencer (Illumina, 100-bp paired-end run mode) at the INRAGeT-PlaGe-GENOTOUL platform. Sequence analyses were performed as previously described by Garcia et al. (2016), using version SL2.50 of the reference tomato genome for read mapping (https://solgenomics.net/ftp//genomes/Solanum_lycopersicum/Heinz1706/assembly/build_2.50/). EMS variants with a read depth between 10 and 100 were considered for allelic frequency analysis. Recombinant BC_1_F_2_ individuals were detected using the EMS-induced SNPs flanking the putative mutation as markers in a Kompetitive allele-specific PCR (KASP) assay (Smith and Maughan, 2015). Subsequent genotyping of the causal mutation was done through Sanger sequencing of PCR products. All primer sequences are listed in **Suppl. Table S1**.

### Generation of CRISPR/Cas9 Lines

CRISPR/Cas9 mutagenesis of *SlLDS1* and tomato cotyledon transformation were performed as described by Bollier et al. (2018), except that the level 1 construct *pICH47742:2×35S-5’UTR-hCas9(STOP)-NOST* was replaced by *pICH47742:2×35S-5’UTR-pcoCas9(STOP)-NOST* (Addgene plasmid # 112079) and the level 1 constructs assembled into the level 2 *pICSL4723* (Addgene plasmid # 86172). The sgRNA target sequence was designed using CRISPR-P 2.0 web software (Lei et al., 2014). The T_0_ plants resulting from the regeneration of the cotyledons were genotyped and their fruits were phenotyped. T_1_ seeds selected from T_0_ plants with rough fruits were sown for further characterization. CRISPR/Cas9 positive lines were further genotyped for indel mutations using primers flanking the target sequence. Resulting mutations were detected by PCR and sequencing in T_0_ and T_1_ lines. List of sgRNA and primers used are given in **Suppl. Table S1**.

### Histological Analyses

Five-millimeter basal sections from shoot explants were fixed in a paraformaldehyde/Triton solution, dehydrated and embedded in Technovit 7,100 resin as previously described (Alaguero-Cordovilla et al., 2021). Sections were stained with 0.05% weight/volume (w/v) toluidine blue (Sigma-Aldrich) and observed using a bright-field Motic BA210 microscope (Motic Spain, Spain). To visualize auxin and CK response, the *DR5:mScarletI-NLS*;*TCSn:mNeonGreen-NLS* reporters were introduced by crossing to MT and *mars1-1* and selected in F_3_ by Sanger sequencing. Hypocotyls from seedlings at the 100–101 growth stages, were cut into 5 mm sections and transferred to Murashige and Skoog medium (MS) basal salt medium with 2% sucrose, 0.8% agar, and 0.30 µM NAA. The expression of the reporter proteins, mScarletI-NLS and mNeonGreen-NLS, were observed at 0, 40, and 75 hae under a ZEISS Axio Observer microscope equipped with the Apotome 2 module (Carl-Zeiss-Stiftung, Oberkochen, Germany). The excitation of mScarletI-NLS was achieved by using a wavelength of 450-490 nm, while the fluorescence emission was collected between 500 and 550 nm. The excitation of mNeonGreen-NLS was achieved by using a wavelength of 538-562 nm, while the fluorescence emission was collected between 570 and 640 nm. Images of the vascular bundles were captured at varying Z-positions of the sample and processed by using ImageJ (Carotenuto et al., 2019).

Histological analyses of pericarp tissue in the fruits were performed as previously described (Musseau et al., 2017, 2020). Transverse fresh sections of fruit exocarp were obtained from three independent 20 DPA stage fruits from MT and *mars1/rough* plants. Transverse sections were stained with calcofluor white stain (Sigma-Aldrich) for cell walls. The sections were submerged for 30 s in 0.05% calcofluor white stain and washed for 5 min with phosphate-buffered saline (PBS) two times. The stained sections were mounted on a slide in fluorescence mounting medium (CitiFluor AF1, England) and observed by confocal microscopy (Zeiss LSM880, Germany). Images were acquired with Zen 2011 software. All these experiments were realized at the Bordeaux Imaging Center (http://www.bic.u-bordeaux.fr/). Observations were made on MT, *mars1/rough* homozygous mutant and wild-type individuals from the mapping BC_1_F_2_ population detected by KASP assay and three CRISPR/Cas9 T_2_ homozygous lines. For each genotype, a minimum of three pericarp sections from three fruits from different plants grown side-by-side were observed.

### RNA Isolation, Next Generation Sequencing and RNA-seq Analysis

In each sample, 3–4 mm of the basal region of the hypocotyl were collected at the indicated times: 0, 8, 20, or 48 hae (**Supp. Fig. 4A**). Three biological replicates, each consisting of 8–10 basal fragments, were harvested and immediately frozen in liquid nitrogen. RNA was extracted as previously described (Larriba et al., 2021a). Total RNA was extracted from ∼100 mg of powdered tissue using the Spectrum Plant Total RNA Kit (Merck, Burlington, MA, USA) and treated with DNAse I (Thermo Fisher Scientific, Waltham, MA, USA); the RNA was then stored at −80 °C. Sequencing libraries and next-generation sequencing were carried out using the BGISEQ-500 pipeline (BGI-Tech, Shenzhen, China). In the first experiment, three libraries were sequenced for the MT and *mars1-3* mutant, and two libraries were sequenced for the P19E6 mutant using stranded DNBSeq protocol (BGI) in Pair End (PE) mode whit 100 cycles of sequencing; in the second experiment, three libraries were sequenced for samples at 8 hae, and two libraries were sequenced for samples at 20 and 48 hae using non-stranded DNBSeq protocol (BGI) in PE with 100 cycles of sequencing (BioProject: PRJNA1289037).

The bioinformatics workflow utilized for RNA-seq analysis is shown in **Supp. Fig. 4B**. Briefly, the sequencing libraries from both experiments were aligned against the tomato SL4.0 genome assembly using the STAR aligner. The assignment of reads to gene and lncRNA models (ITAG4.0 annotation) was conducted with featureCounts tool from the Subread package. The number of reads was determined using the FeatureCounts tool with the -s 2 parameters in Experiment 1 and the default setting in Experiment 2. The reads were normalized using counts per million (CPM), and differential expression analysis was performed using DESeq2, from iDEP 2.0 (Ge et al., 2018) to identify differentially expressed genes with a false discovery rate (FDR) = 0.01 and fold change (FC) = |1.5|. The expressed transcripts (X_LOC) were obtained in the collapsed BAM files of each time/condition and genotype of Experiment 1 using Stringtie and GFFCompare. The number of reads associated with the identified X_LOC in the collapsed BAMs of both experiments, differentiated by time and genotype, were quantified using the FeatureCounts tool from the GFF. To calculate the FC of each X_LOC, we utilized the counts values per genotype, calculating the differential between CPM values between the mutant and control genotypes. The chromosomal positions of lncRNAs were annotated in relation to the genes annotated in ITAG4 using GFFCompare.

### Protein Extraction and Western Blot Analysis

Nucleus isolation was performed as described by Tian et al. (2020). Briefly, 1.5 g of frozen leaf tissue from four-week-old MT and *mars1-3* plants was homogenized in liquid nitrogen. The tissue was incubated in 15 mL of nucleus isolation buffer (NIB) for 30 minutes, inverted every five minutes. The mixture was filtered sequentially through 70- and 40-µm filters. The filtrate was centrifuged at 900 g for 10 minutes at 4 °C. The resulting pellet containing the nuclei was washed with 1 mL of NIB and then centrifuged again at 900 g for 10 minutes. Histones were isolated from the nuclei using the acid extraction protocol described by Shechter et al. (2007). One mL of 4 N HLSOL was added to the pellet, incubated for one hour with rotation at 4 °C, and was then centrifuged at 21,000 g for 10 minutes at 4 °C to obtain the histone-containing supernatant. Histone precipitation was carried out with trichloroacetic acid at a final concentration of 33% for one hour at 4 °C. The histones were then precipitated by centrifugation, washed twice with acetone, dried, and resuspended in nuclease-free water. The proteins were separated on a 15% SDS-polyacrylamide gel and, subsequently, blotted onto PVDF blotting membranes (GE HealthCare Technologies, Chicago, IL, USA) as indicated elsewhere (Kenesi et al., 2023). The PVDF membrane was immersed in the blocking solution overnight. The blocking solution used was TBS-T (150 mM NaCl, 50 mM Tris-HCl, 0.2% Tween; pH 7.5) with 5% non-fat milk (Kenesi et al., 2023). Membranes were subsequently washed with TBS-T at least three times. After washing the membranes, they were incubated with histone-specific primary antibodies (anti-H3K4me1 [Abcam, ab1791], anti-H3K4me2 [Abcam, ab176882], anti-H3K4me3 [Agrisera, AS16 3190], anti-H3K36me3 [Abcam, ab9050], or anti-acetyl Lys-9 [Sigma Aldrich 07-352],) for 4 hours at room temperature. The membranes were then washed with TBS-T and then incubated for 1 hour with secondary antibodies (Goat-Anti-Rabbit-HRP: 1:10000 [Agrisera, AS09602], or Goat-Anti-mouse-HRP: 1:5000 [GE HealthCare, NA 931VS]) at room temperature. Immunoreactive signal was subsequently developed with the use of WesternSure PREMIUM chemiluminescent substrate and detected by using the ChemiDoc XRS+ imaging system (Bio-Rad, Hercules, CA, USA).

### Reverse Transcription, PCR, D-Topo Cloning and Sequencing

Total RNA was extracted from approximately 80-100 mg of shoot explants, mature leaves, and ripe green and red fruits using TRIzol reagent (Invitrogen, Carlsbad, CA, USA) according to the manufacturer’s protocols. The RNA obtained was subjected to retrotranscription (RT) using the RevertAid kit (ThermoFisher Scientific, Waltham, MA, USA) according to the manufacturer’s recommendations. The resulting cDNA was used as a template for PCR amplification using several primer pairs listed as RT-PCR in **Supp. Table S1**. PCR products amplified with Vacuolar2F_DT and CYC-3R primers (**Suppl. Table S1**) were cloned using the pENTR/D-TOPO kit (ThermoFisher Scientific) and unique clones, as determined by the size of PCR products using M13F and M13R primers, were Sanger sequenced. For sequencing the *Solyc04g081650-Solyc04g081670* genomic region, PCR products were amplified using the primers listed in **Supp. Table S1** as DNA sequencing.

### Vector Construction and Agroinjection for Virus Induced Gene Silencing (VIGS)

The SGN VIGS Tool (Fernandez-Pozo et al., 2015; https://vigs.solgenomics.net/), was used to design specific probes against *Solyc04g081650* and *Solyc04g081660* within the gene supercluster, and against *Solyc03g123760*, which encodes the phytoene desaturase that was used as a positive control. Primers listed as VIGS in **Supp. Table S1** were used for PCR amplification. PCR products were cloned into the pTRV2-GG vector (plasmid #105349, Addgene) using Eco31I, and positive clones were verified by Sanger sequencing. pTRV1 and pTRV2-GG vectors were electroporated into *Agrobacterium tumefaciens* GV3101 strains and selected based on antibiotic resistance and PCR. *A. tumefaciens* strains containing pTRV1 or pTRV2-GG (either empty or with *Solyc04g081650*, *Solyc04g081660* or *Solyc03g123760* probes) were grown in YEB medium plus antibiotics for 2 days at 28°C, cells were harvested by centrifugation afterwards and resuspended in the infiltration buffer (10 mM MgCl_2_, 10 mM MES pH 5.6, 150 mM acetosyringone) to a final OD600 of 0.5 in a 1:1 ratio. This infiltration solution was injected through the carpopodium of young tomato fruits (5 to 8 DPA) attached to the plant using a 1 ml syringe. For each pTRV2-GG strain, three replicates of the agroinjection protocol were performed, each with a minimum of 3-5 tomatoes. The injected fruits were observed every week for one month.

### Statistical Analyses

The descriptive statistics were calculated using StatGraphics Centurion 16.1.03 (StatPoint Technologies, Inc. Warrenton, VA, USA) and GraphPad Prism version 8.3.1 for Windows (GraphPad Software, La Jolla California USA) software. We performed multiple testing analyses using the ANOVA F-test or the Fisher’s least significant difference (LSD) methods (p–value<0.01, unless otherwise indicated). Nonparametric tests were used when necessary.

## Acknowledgements

We thank Dr. Idan Efroni (The Hebrew University, Israel) for sharing seeds from the *DR5:mScarletI-NLS*;*TCSn:mNeonGreen-NLS* reporter line, Prof. Johannes Stuttmann (Institute of Biosciences and Biotechnologies, France) for providing pTRV2-GG and PTRV1 plasmids, and Profs. José Luis Micol, María Rosa Ponce, Miguel Ángel Sogorb, and Eduardo Fernández (Instituto de Bioingeniería, UMH, Spain) for sharing their equipment. This research is supported by the grant PID2021-126840OB-I00 funded by MCIN/AEI/10.13039/501100011033, and by the “ERDF A way of making Europe”. RR and LC hold a postdoctoral contract (CIAPOS/2023/138) and “Programa INVESTIGO” contract (INVEST/2022/247) from the Generalitat Valenciana.

## Author contribution statement

EL, CB, AA-C, JP, RR, J-PM, LC, BB and D.E.-B. designed and performed all the experiments. The first draft of the manuscript was written by CR and JMP-P. JMP-P wrote the final manuscript. All authors read and approved the final manuscript.

## Availability of data and materials

All data generated or analyzed during this study are provided in this article and its supplementary data files or it will be provided upon a reasonable request.

## References

Alaguero-Cordovilla, A., Gran-Gómez, F. J., Jadczak, P., Mhimdi, M., Ibáñez, S., Bres, C., Just, D., Rothan, C., and Pérez-Pérez, J. M. (2020). A quick protocol for the identification and characterization of early growth mutants in tomato. Plant Sci. 301:110673.

Alaguero-Cordovilla, A., Sánchez-García, A. B., Ibáñez, S., Albacete, A., Cano, A., Acosta, M., and Pérez-Pérez, J. M. (2021). An auxin-mediated regulatory framework for wound-induced adventitious root formation in tomato shoot explants. Plant Cell Environ. Advance Access published 2021, doi:10.1111/pce.14001.

An, J., Chaouche, R. B., Pereyra-Bistraín, L. I., Zalzalé, H., Wang, Q., Huang, Y., He, X., Lopes, C. D., Antunez-Sanchez, J., Bergounioux, C., et al. (2024). An atlas of the tomato epigenome reveals that KRYPTONITE shapes TAD-like boundaries through the control of H3K9ac distribution. Proc. Natl. Acad. Sci. U. S. A. 121:e2400737121.

Berr, A., Shafiq, S., and Shen, W. H. (2011). Histone modifications in transcriptional activation during plant development. Biochim. Biophys. Acta - Gene Regul. Mech. 1809:567–576.

Bollier, N., Sicard, A., Leblond, J., Latrasse, D., Gonzalez, N., Gévaudant, F., Benhamed, M., Raynaud, C., Lenhard, M., Chevalier, C., et al. (2018). At-MINI ZINC FINGER2 and Sl-INHIBITOR OF MERISTEM ACTIVITY, a Conserved Missing Link in the Regulation of Floral Meristem Termination in Arabidopsis and Tomato. Plant Cell 30:83.

Caro, E., Stroud, H., Greenberg, M. V. C., Bernatavichute, Y. V, Feng, S., Groth, M., Vashisht, A. A., Wohlschlegel, J., and Jacobsen, S. E. (2012). The SET-domain protein SUVR5 mediates H3K9me2 deposition and silencing at stimulus response genes in a DNA methylation–independent manner Advance Access published 2012.

Carotenuto, G., Sciascia, I., Oddi, L., Volpe, V., and Genre, A. (2019). Size matters: Three methods for estimating nuclear size in mycorrhizal roots of Medicago truncatula by image analysis. BMC Plant Biol. 19:1–14.

Czerednik, A., Busscher, M., Bielen, B. A. M., Wolters-Arts, M., De Maagd, R. A., and Angenent, G. C. (2012). Regulation of tomato fruit pericarp development by an interplay between CDKB and CDKA1 cell cycle genes. J. Exp. Bot. 63:2605–2617.

Da, G., Lenkart, J., Zhao, K., Shiekhattar, R., Cairns, B. R., and Marmorstein, R. (2006). Structure and function of the SWIRM domain, a conserved protein module found in chromatin regulatory complexes. Proc. Natl. Acad. Sci. U. S. A. 103:2057–2062.

Dahan-Meir, T., Filler-Hayut, S., Melamed-Bessudo, C., Bocobza, S., Czosnek, H., Aharoni, A., and Levy, A. A. (2018). Efficient in planta gene targeting in tomato using geminiviral replicons and the CRISPR/Cas9 system. Plant J. 95:5–16.

Davey, N. E., and Morgan, D. O. (2016). Building a Regulatory Network with Short Linear Sequence Motifs: Lessons from the Degrons of the Anaphase-Promoting Complex. Mol. Cell 64:12–23.

Dewitte, W., Riou-Khamlichi, C., Scofield, S., Healy, J. M. S., Jacqmard, A., Kilby, N. J., and Murray, J. A. H. (2003). Altered Cell Cycle Distribution, Hyperplasia, and Inhibited Differentiation in Arabidopsis Caused by the D-Type Cyclin CYCD3. Plant Cell 15:79–92.

Du, J., Johnson, L. M., Groth, M., Feng, S., Hale, C. J., Li, S., Vashisht, A. A., Gallego-Bartolome, J., Wohlschlegel, J. A., Patel, D. J., et al. (2014). Mechanism of DNA methylation-directed histone methylation by KRYPTONITE. Mol. Cell 55:495–504.

Eloy, N. B., De Freitas Lima, M., Van Damme, D., Vanhaeren, H., Gonzalez, N., De Milde, L., Hemerly, A. S., Beemster, G. T. S., Inzé, D., and Ferreira, P. C. G. (2011). The APC/C subunit 10 plays an essential role in cell proliferation during leaf development. Plant J. 68:351–363.

Fernandez-Pozo, N., Rosli, H. G., Martin, G. B., and Mueller, L. A. (2015). The SGN VIGS tool: User-friendly software to design virus-induced gene silencing (VIGS) Constructs for functional genomics. Mol. Plant 8:486–488.

Fu, D. Q., Zhu, B. Z., Zhu, H. L., Jiang, W. B., and Luo, Y. B. (2005). Virus-induced gene silencing in tomato fruit. Plant J. 43:299–308.

Gan, L., Song, M., Wang, X., Yang, N., Li, H., Liu, X., and Li, Y. (2022). Cytokinins are involved in regulation of tomato pericarp thickness and fruit size. Hortic. Res. 9.

Gao, Y., Yang, S., Yuan, L., Cui, Y., and Wu, K. (2012). Comparative Analysis of SWIRM Domain-Containing Proteins in Plants. Int. J. Genomics 2012:310402.

Garcia, V., Bres, C., Just, D., Fernandez, L., Tai, F. W. J., Mauxion, J. P., Le Paslier, M. C., Bérard, A., Brunel, D., Aoki, K., et al. (2016). Rapid identification of causal mutations in tomato EMS populations via mapping-by-sequencing. Nat. Protoc. 11:2401–2418.

Ge, S. X., Son, E. W., and Yao, R. (2018). iDEP: an integrated web application for differential expression and pathway analysis of RNA-Seq data. BMC Bioinforma. 2018 191 19:1–24.

Gonzalez, N., Gévaudant, F., Hernould, M., Chevalier, C., and Mouras, A. (2007). The cell cycle-associated protein kinase WEE1 regulates cell size in relation to endoreduplication in developing tomato fruit. Plant J. 51:642–655.

Greenberg, M. V. C., Deleris, A., Hale, C. J., Liu, A., Feng, S., and Jacobsen, S. E. (2013). Interplay between Active Chromatin Marks and RNA-Directed DNA Methylation in Arabidopsis thaliana. PLOS Genet. 9:e1003946.

Hosmani, P. S., Flores-Gonzalez, M., Geest, H. van de, Maumus, F., Bakker, L. V., Schijlen, E., Haarst, J. van, Cordewener, J., Sanchez-Perez, G., Peters, S., et al. (2019). An improved de novo assembly and annotation of the tomato reference genome using single-molecule sequencing, Hi-C proximity ligation and optical maps. bioRxiv Advance Access published September 14, 2019, doi:10.1101/767764.

Hsieh, J.-W. A., Yen, M.-R., Hung, F.-Y., Wu, K., and Chen, P.-Y. (2024). Epigenetic factors direct synergistic and antagonistic regulation of transposable elements in Arabidopsis. Plant Physiol. 196:1939–1952.

Hung, F. Y., Chen, F. F., Li, C., Chen, C., Lai, Y. C., Chen, J. H., Cui, Y., and Wu, K. (2018). The Arabidopsis LDL1/2-HDA6 histone modification complex is functionally associated with CCA1/LHY in regulation of circadian clock genes. Nucleic Acids Res. 46:10669–10681.

Hung, F. Y., Chen, C., Yen, M. R., Hsieh, J. W. A., Li, C., Shih, Y. H., Chen, F. F., Chen, P. Y., Cui, Y., and Wu, K. (2020). The expression of long non-coding RNAs is associated with H3Ac and H3K4me2 changes regulated by the HDA6-LDL1/2 histone modification complex in Arabidopsis. NAR Genomics Bioinforma. 2.

Ikeuchi, M., Ogawa, Y., Iwase, A., and Sugimoto, K. (2016). Plant regeneration: cellular origins and molecular mechanisms. Development 143:1442–1451.

Ikeuchi, M., Iwase, A., Rymen, B., Lambolez, A., Kojima, M., Takebayashi, Y., Heyman, J., Watanabe, S., Seo, M., De Veylder, L., et al. (2017). Wounding triggers callus formation via dynamic hormonal and transcriptional changes. Plant Physiol. 175:1158–1174.

Ikeuchi, M., Favero, D. S., Sakamoto, Y., Iwase, A., Coleman, D., Rymen, B., and Sugimoto, K. (2019). Molecular Mechanisms of Plant Regeneration. Annu. Rev. Plant Biol. 70:377–406.

Inagaki, S., Takahashi, M., Hosaka, A., Ito, T., Toyoda, A., Fujiyama, A., Tarutani, Y., and Kakutani, T. (2017). GeneLbody chromatin modification dynamics mediate epigenome differentiation in Arabidopsis. EMBO J. 36:970–980.

Isaacson, T., Kosma, D. K., Matas, A. J., Buda, G. J., He, Y., Yu, B., Pravitasari, A., Batteas, J. D., Stark, R. E., Jenks, M. A., et al. (2009). Cutin deficiency in the tomato fruit cuticle consistently affects resistance to microbial infection and biomechanical properties, but not transpirational water loss. Plant J. 60:363–377.

Iwase, A., Harashima, H., Ikeuchi, M., Rymen, B., Ohnuma, M., Komaki, S., Morohashi, K., Kurata, T., Nakata, M., Ohme-Takagi, M., et al. (2017). WIND1 promotes shoot regeneration through transcriptional activation of ENHANCER OF SHOOT REGENERATION1 in arabidopsis. Plant Cell 29:54–69.

Jumper, J., Evans, R., Pritzel, A., Green, T., Figurnov, M., Ronneberger, O., Tunyasuvunakool, K., Bates, R., Žídek, A., Potapenko, A., et al. (2021). Highly accurate protein structure prediction with AlphaFold. Nat. 2021 5967873 596:583–589.

Just, D., Garcia, V., Fernandez, L., Bres, C., Mauxion, J.-P., Petit, J., Jorly, J., Assali, J., Ceacute, Bournonville, L., et al. (2013). Micro-Tom mutants for functional analysis of target genes and discovery of new alleles in tomato. Plant Biotechnol. 30:225–231.

Kenesi, E., Kolbert, Z., Kaszler, N., Klement, É., Ménesi, D., Molnár, Á., Valkai, I., Feigl, G., Rigó, G., Cséplő, Á., et al. (2023). The ROP2 GTPase Participates in Nitric Oxide (NO)-Induced Root Shortening in Arabidopsis. Plants 12:750.

Larriba, E., Sánchez-García, A. B., Martínez-Andújar, C., Albacete, A., and Pérez-Pérez, J. M. (2021a). Tissue-specific metabolic reprogramming during wound-induced organ formation in tomato hypocotyl explants. Int. J. Mol. Sci. 22:10112.

Larriba, E., Belén Sánchez-García, A., Salud Justamante, M., Martínez-Andújar, C., Albacete, A., Manuel Pérez-Pérez, J., Tran, P., and Golam Mostofa, M. (2021b). Dynamic Hormone Gradients Regulate Wound-Induced de novo Organ Formation in Tomato Hypocotyl Explants. Int. J. Mol. Sci. 2021, Vol. 22, Page 11843 22:11843.

Larriba, E., Nicolás-Albujer, M., Sánchez-García, A. B., and Pérez-Pérez, J. M. (2022). Identification of Transcriptional Networks Involved in De Novo Organ Formation in Tomato Hypocotyl Explants. Int. J. Mol. Sci. 23:16112.

Lashbrooke, J., Cohen, H., Levy-Samocha, D., Tzfadia, O., Panizel, I., Zeisler, V., Massalha, H., Stern, A., Trainotti, L., Schreiber, L., et al. (2016). MYB107 and MYB9 Homologs Regulate Suberin Deposition in Angiosperms. Plant Cell 28:2097–2116.

Lee, K., Park, O. S., and Seo, P. J. (2017). Arabidopsis ATXR2 deposits H3K36me3 at the promoters of LBD genes to facilitate cellular dedifferentiation. Sci. Signal. 10.

Lei, Y., Lu, L., Liu, H. Y., Li, S., Xing, F., and Chen, L. L. (2014). CRISPR-P: A web tool for synthetic single-guide RNA design of CRISPR-system in plants. Mol. Plant 7:1494–1496.

Ma, J., Li, Q., Zhang, L., Cai, S., Liu, Y., Lin, J., Huang, R., Yu, Y., Wen, M., and Xu, T. (2022). High auxin stimulates callus through SDG8-mediated histone H3K36 methylation in Arabidopsis. J. Integr. Plant Biol. 64:2425–2437.

Mathew, M. M., and Prasad, K. (2021). Model systems for regeneration: Arabidopsis. Dev. 148.

Metzger, E., Wissmann, M., Yin, N., Müller, J. M., Schneider, R., Peters, A. H. F. M., Günther, T., Buettner, R., and Schüle, R. (2005). LSD1 demethylates repressive histone marks to promote androgen-receptor-dependent transcription. Nat. 2005 4377057 437:436–439.

Mozgova, I., Wildhaber, T., Trejo-Arellano, M. S., Fajkus, J., Roszak, P., Köhler, C., and Hennig, L. (2018). Transgenerational phenotype aggravation in CAF-1 mutants reveals parent-of-origin specific epigenetic inheritance. New Phytol. 220:908–921.

Musseau, C., Just, D., Jorly, J., Gévaudant, F., Moing, A., Chevalier, C., Lemaire-Chamley, M., Rothan, C., and Fernandez, L. (2017). Identification of two new mechanisms that regulate fruit growth by cell expansion in tomato. Front. Plant Sci. 8:259220.

Musseau, C., Jorly, J., Gadin, S., Sørensen, I., Deborde, C., Bernillon, S., Mauxion, J. P., Atienza, I., Moing, A., Lemaire-Chamley, M., et al. (2020). The Tomato Guanylate-Binding Protein SlGBP1 Enables Fruit Tissue Differentiation by Maintaining Endopolyploid Cells in a Non-Proliferative State. Plant Cell 32:3188–3205.

Nadakuduti, S. S., Pollard, M., Kosma, D. K., Allen, C., Ohlrogge, J. B., and Barry, C. S. (2012). Pleiotropic Phenotypes of the sticky peel Mutant Provide New Insight into the Role of CUTIN DEFICIENT2 in Epidermal Cell Function in Tomato. Plant Physiol. 159:945–960.

Nafati, M., Frangne, N., Hernould, M., Chevalier, C., and Gévaudant, F. (2010). Functional characterization of the tomato cyclin-dependent kinase inhibitor SlKRP1 domains involved in protein-protein interactions. New Phytol. 188:136–149.

Omary, M., Gil-Yarom, N., Yahav, C., Steiner, E., Hendelman, A., and Efroni, I. (2022). A conserved superlocus regulates above- and belowground root initiation. Science 375:eabf4368.

Pattison, R. J., and Catalá, C. (2012). Evaluating auxin distribution in tomato (Solanum lycopersicum) through an analysis of the PIN and AUX/LAX gene families. Plant J. 70:585–598.

Petit, J., Bres, C., Mauxion, J. P., Tai, F. W. J., Martin, L. B. B., Fich, E. A., Joubès, J., Rose, J. K. C., Domergue, F., and Rothan, C. (2016). The Glycerol-3-Phosphate Acyltransferase GPAT6 from Tomato Plays a Central Role in Fruit Cutin Biosynthesis. Plant Physiol. 171:894–913.

Petit, J., Bres, C., Reynoud, N., Lahaye, M., Marion, D., Bakan, B., and Rothan, C. (2021). Unraveling Cuticle Formation, Structure, and Properties by Using Tomato Genetic Diversity. Front. Plant Sci. 12:778131.

Renaudin, J. P., Deluche, C., Cheniclet, C., Chevalier, C., and Frangne, N. (2017). Cell layer-specific patterns of cell division and cell expansion during fruit set and fruit growth in tomato pericarp. J. Exp. Bot. 68:1613–1623.

Renaudin, J. P., Cheniclet, C., Rouyère, V., Chevalier, C., and Frangne, N. (2023). The Cell Pattern of Tomato Fruit Pericarp is Quantitatively and Differentially Regulated by the Level of Gibberellin in Four Cultivars. J. Plant Growth Regul. 42:5945–5958.

Reynoud, N., Geneix, N., D’Orlando, A., Petit, J., Mathurin, J., Deniset-Besseau, A., Marion, D., Rothan, C., Lahaye, M., and Bakan, B. (2023). Cuticle architecture and mechanical properties: a functional relationship delineated through correlated multimodal imaging. New Phytol. 238:2033–2046.

Rothan, C., Just, D., Fernandez, L., Atienza, I., Ballias, P., and Lemaire-Chamley, M. (2016). Culture of the tomato Micro-Tom Cultivar in greenhouse. In Methods in Molecular Biology, pp. 57–64. Humana Press Inc.

Roudier, F., Ahmed, I., Bérard, C., Sarazin, A., Mary-Huard, T., Cortijo, S., Bouyer, D., Caillieux, E., Duvernois-Berthet, E., Al-Shikhley, L., et al. (2011). Integrative epigenomic mapping defines four main chromatin states in Arabidopsis. EMBO J. 30:1928–1938.

Sakamoto, Y., Kawamura, A., Suzuki, T., Segami, S., Maeshima, M., Polyn, S., De Veylder, L., and Sugimoto, K. (2022). Transcriptional activation of auxin biosynthesis drives developmental reprogramming of differentiated cells. Plant Cell 34:4348–4365.

Shechter, D., Dormann, H.L., Allis, C.D., and Hake, S.B. (2007). Extraction, purification and analysis of histones. Nat. Protoc. 2:1445–1457.

Shi, Y., Lan, F., Matson, C., Mulligan, P., Whetstine, J. R., Cole, P. A., Casero, R. A., and Shi, Y. (2004). Histone demethylation mediated by the nuclear amine oxidase homolog LSD1. Cell 119:941–953.

Shikata, M., and Ezura, H. (2016). Micro-tom tomato as an alternative plant model system: Mutant collection and efficient transformation. In Methods in Molecular Biology, pp. 47–55. Humana Press Inc.

Shim, S., Lee, H. G., Park, O. S., Shin, H., Lee, K., Lee, H., Huh, J. H., and Seo, P. J. (2022). Dynamic changes in DNA methylation occur in TE regions and affect cell proliferation during leaf-to-callus transition in Arabidopsis. Epigenetics 17:41–58.

Smith, S. M., and Maughan, P. J. (2015). SNP genotyping using KASPar assays. Methods Mol. Biol. 1245:243–256.

Tian, C., Du, Q., Xu, M., Du, F., and Jiao, Y. (2020). Single-nucleus RNA-seq resolves spatiotemporal developmental trajectories in the tomato shoot apex. bioRxiv [preprint] 2020.09.20.305029.

Tourdot, E., Martin, P. G. P., Maza, E., Mauxion, J. P., Djari, A., Gévaudant, F., Chevalier, C., Pirrello, J., and Gonzalez, N. (2024). Ploidy-specific transcriptomes shed light on the heterogeneous identity and metabolism of developing tomato pericarp cells. Plant J. 118:997–1015.

Vodermaier, H. C. (2004). APC/C and SCF: Controlling Each Other and the Cell Cycle. Curr. Biol. 14:R787–R796.

Weingartner, M., Criqui, M. C., Mészáros, T., Binarova, P., Schmit, A. C., Helfer, A., Derevier, A., Erhardt, M., Bögre, L., and Genschik, P. (2004). Expression of a Nondegradable Cyclin B1 Affects Plant Development and Leads to Endomitosis by Inhibiting the Formation of a Phragmoplast. Plant Cell 16:643–657.

Xu, L., and Huang, H. (2014). Genetic and Epigenetic Controls of Plant Regeneration. In Current Topics in Developmental Biology, pp. 1–33. Academic Press.

Yang, W., Zhai, H., Wu, F., Deng, L., Chao, Y., Meng, X., Chen, Q., Liu, C., Bie, X., Sun, C., et al. (2024). Peptide REF1 is a local wound signal promoting plant regeneration. Cell 187:3024–3038.e14.

Zhang, X., Bernatavichute, Y. V., Cokus, S., Pellegrini, M., and Jacobsen, S. E. (2009). Genome-wide analysis of mono-, di- and trimethylation of histone H3 lysine 4 in Arabidopsis thaliana. Genome Biol. 10:1–14.

Zhang, G., Zhao, F., Chen, L., Pan, Y., Sun, L., Bao, N., Zhang, T., Cui, C. X., Qiu, Z., Zhang, Y., et al. (2019). Jasmonate-mediated wound signalling promotes plant regeneration. Nat. Plants 5:491–497.

