## Supplementary material for "Role of the tomato *MARS1/ROUGH* gene encoding a LYSINE-SPECIFIC HISTONE DEMETHYLASE 1 in adventitious root and fruit skin formation": Supp. Fig

### Suppl. Figure S1

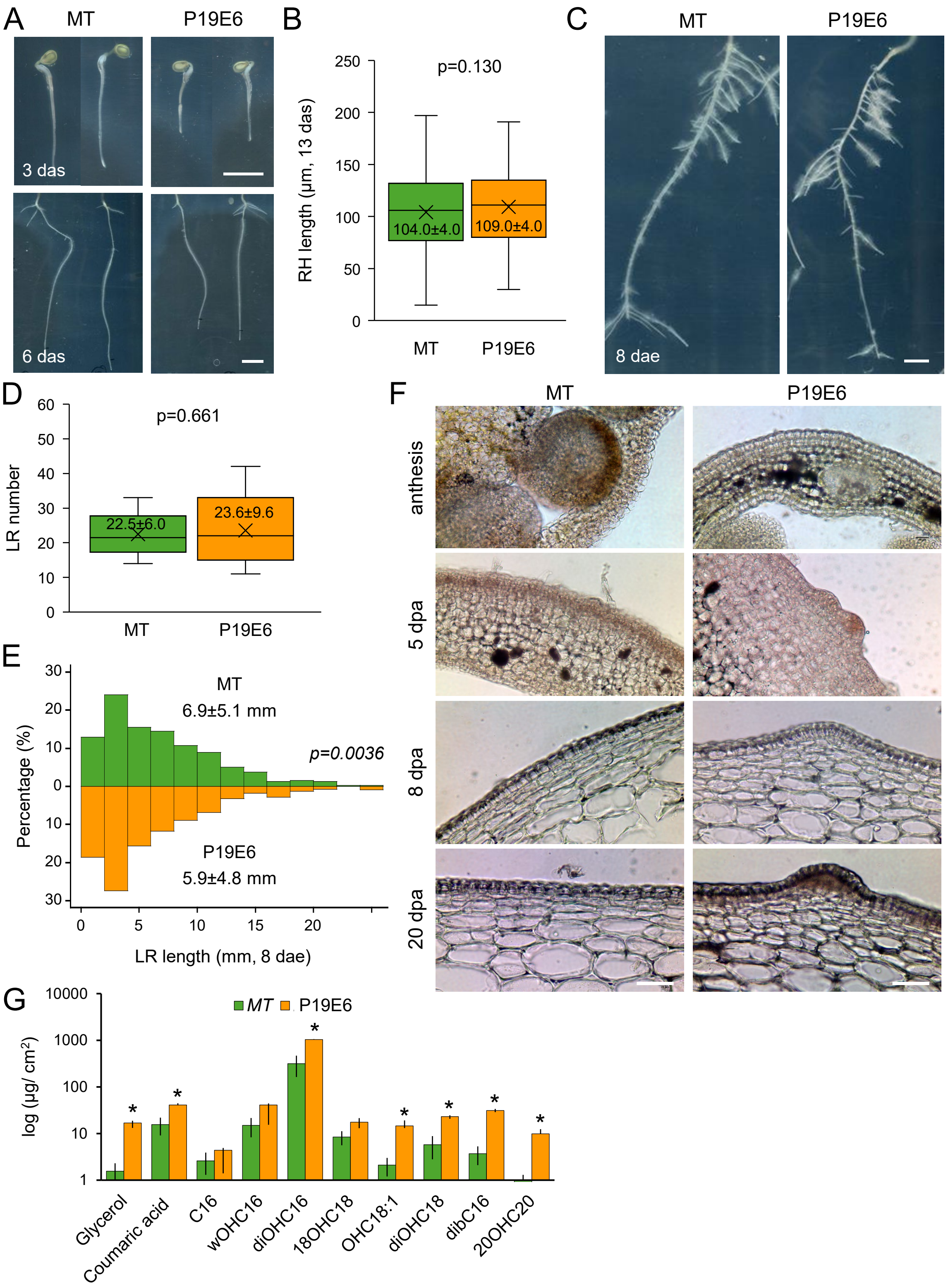


**Suppl. Figure S1. Root phenotype in the P19E6 mutant.** (A) Representative images of young seedlings at 3 and 6 days after sowing (das). (B) Box plot representation of root hair (RH) length in the differentiated region of the PR. (C) Representative images of the entire root system of selected genotypes at 8 days after PR tip excision (dae). (D) Box plot representation of LR numbers at 8 days after PR tip excision. (E) The distribution of LR lengths in MT (top, green) and P19E6 (bottom, orange) mutants at 8 days after PR tip excision. The numbers in B, D, and E indicate the mean ± standard deviation. The p-values are indicated, with those in italics indicating statistical significance (T-test; p-value<0.05; n>21 [D], n>200 [B, E]). (F) Light microscopy of fruit pericarp in WT and P19E6 mutant; dpa, day post anthesis. (G) Monomeric composition of cutin. C16, hexadecanoic acid; wOHC16, 16-hydroxyhexadecanoic acid; diOHC16, (9)10, 16-dihydroxyhexadecanoic acid; 18OHC18, 18-hydroxyoctadecanoic acid; 18OHC18:1, 18-hydroxyoctadecenoic acid; diOHC18, 10, 18-dihydroxyoctadecanoic acid; dibC16, hexadecan-1,16-dioic acid; 20OHC20, 20-hydroxyeicosanoic acid. Asterisks indicate significant differences between P19E6 and MT cutin (student test p<0.05). Scale bars: 2 mm (A, C), and 50 µm (F).

### Suppl. Figure S2

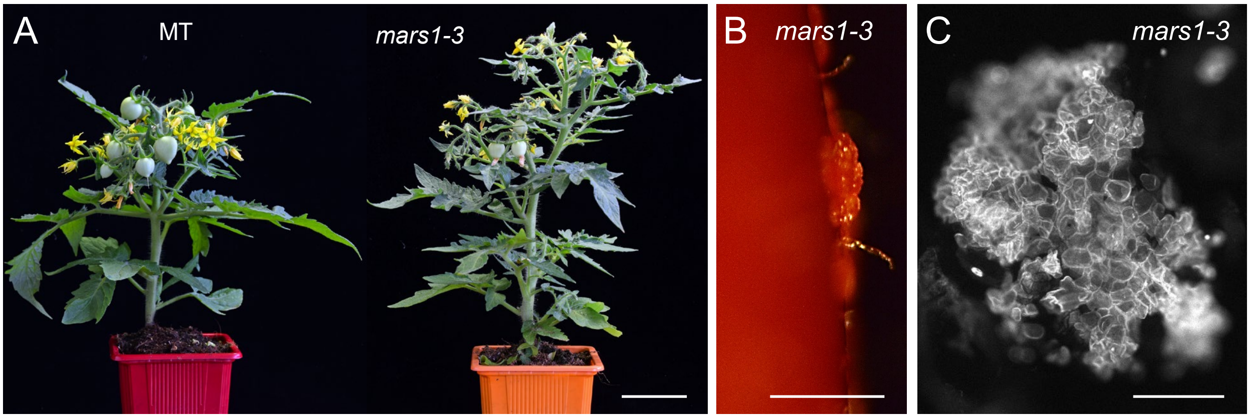


**Supp. Figure S2. Pleiotropic phenotype in *mars1/rough* mutants.** (A) Representative images of the vegetative phenotype of adult plants after 5 weeks in the growth chamber. (B) Details of the fruit surface displaying tissue outgrowth in the *mars1/rough* mutants. (C) Microscopic observation of the tissue outgrowths shown in B. Scale bars: 2 cm (A), 1 mm (B), and 300 µm (C).

### Suppl. Figure S3

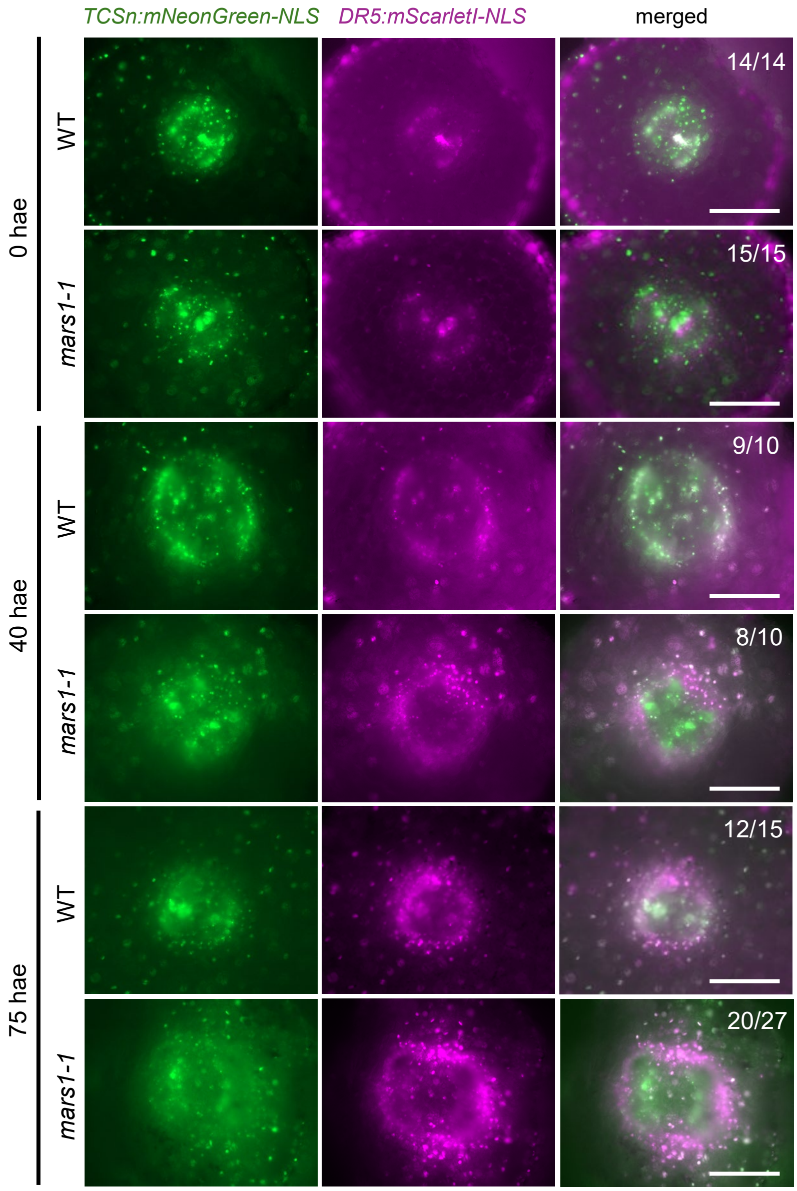


**Supp. Figure S3. Auxin and cytokinin response in the vascular cylinder of *mars1/rough* mutants during AR formation.** Representative images of the basal region of the hypocotyl during AR formation in WT and *mars1-1* lines expressing *DR5:mScarletI-NLS* and *TCSn:mNeonGreen-NLS* markers (Omary et al., 2022) during AR development. Hypocotyl sections were incubated in growth medium supplemented with 0.3 µM IAA at different time points (0, 40, and 75 hours after excision; hae). Scale bars: 250 µm.

### Suppl. Figure S4

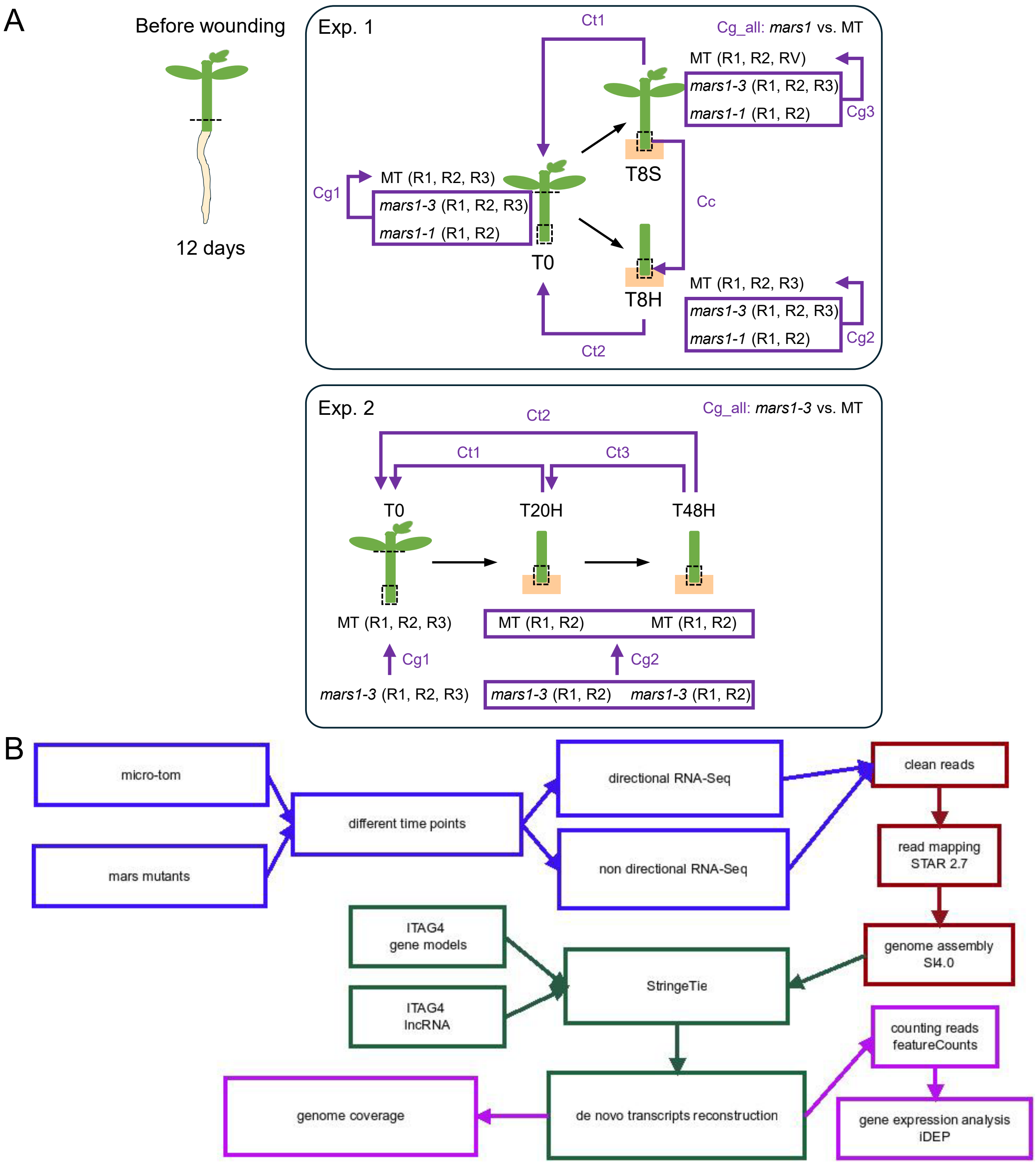


**Supp. Figure S4. Experimental design used for the RNA-seq experiments.** The whole root systems of 12-days-old tomato seedlings were excised, and 4-6 mm of the basal region of the hypocotyl were collected afterwards (T0). In the first experiment (Exp 1), the shoot apex was excised in half of the samples, and the basal regions of the hypocotyl were collected 8 hae (T8H and T8S). In the second experiment, the shoot apex was excised in all the samples and the basal regions of the hypocotyl were collected 20 hae (T20H) and 48 hae (T48H). The number of replicates (R) used are indicated in each case; Rv: virtual replicate. The contrasts implemented for DEG analysis are shown by purple lines with arrows, considering different conditions (Cc), genotypes (Cg) and time points (Ct). In the Cg_all contrast, we compared all libraries from *mars/rough* mutants vs. MT in each experiment. (B) The bioinformatics pipeline employed in our RNA-seq analyses is illustrated in the accompanying flow charts. The steps pertaining to sample selection and the type of RNA-Seq are represented in blue, while all mapping steps are indicated in red. The de novo RNA reconstruction steps are denoted in green, and the final steps of the analysis are highlighted in lilac.

### Suppl. Figure S5

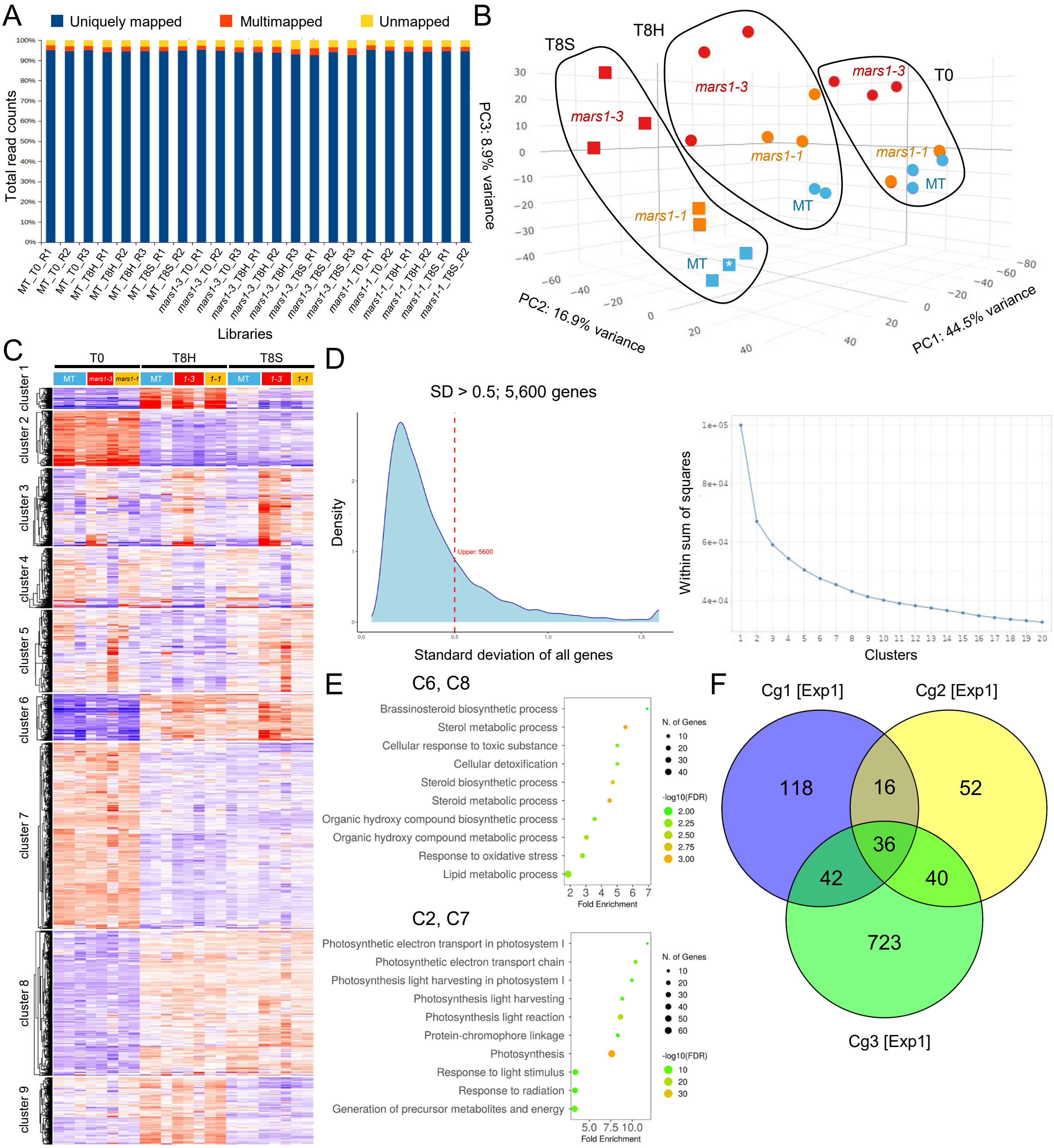


**Supp. Figure S5. Read mapping and analysis of gene expression in experiment 1.** (A) Mapping results of the 23 sequencing libraries from this experiment. (B) Three-dimensional analysis of principal components (PCs) of the RNA-seq results. Samples from the same time point are enclosed with lines of the same color. Note that a virtual replicate (RV) has been used in WT T8S (white asterisk). (C) Clustering of 5,600 genes and lncRNA with standard deviation (SD)>0.5. Predicted clusters are based on time (0, 8 hae) and condition (hypocotyl [H] and shoot [S] explants). Expression values are adjusted on a scale of −4 (blue) to +4 (red). (D) Distribution of SD of 23,784 expressed genes and k-means elbow plot for cluster number estimation. (E) GO BP term-enrichment of the upregulated (clusters 6 and 8), and downregulated (clusters 2 and 7) genes from the k-means clustering. (F) Venn diagrams of DEGs from contrasts comparing *mars1* mutants with MT, as indicated in the main text and Suppl. Fig. S4A. Numbers indicate the DEGs found in each contrast (p-value<0.01).

### Suppl. Figure S6

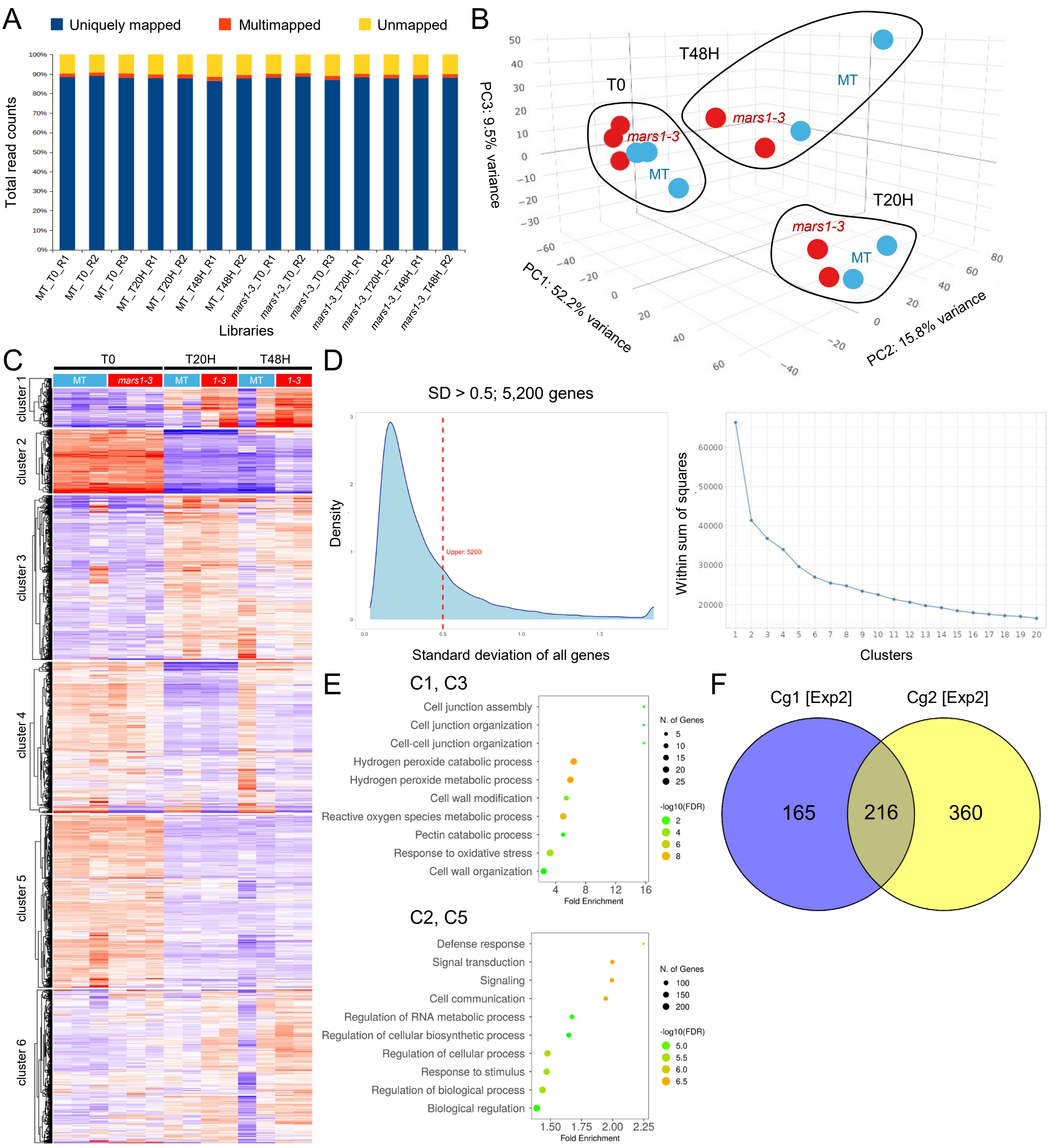


**Supp. Figure S6. Read mapping and analysis of gene expression in experiment 2.** (A) Mapping results of the 14 sequencing libraries. (B) Three-dimensional analysis of principal components (PCs) of the RNA-seq results. Samples from the same time point are enclosed with dark lines. (C) Clustering of 5,200 genes with standard deviation (SD)>0.5. Predicted clusters are based on temporal expression profile ranging from 0 (T0) to 48 (T48H) hae. Expression values are adjusted on a scale of −4 (blue) to +4 (red). (D) Distribution of SD of 22,765 expressed genes and k-means elbow plot for cluster number estimation. (E) GO term-enrichment of the upregulated (clusters 1 and 3), and upregulated genes (clusters 2 and 5) from the k-means clustering. (F) Venn diagrams of DEGs from contrasts comparing the *mars1-3* mutants with MT, as indicated in the main text and Suppl. Fig. S4A. Numbers indicate the DEGs found in each contrast (p-value<0.01).

### Suppl. Figure S7

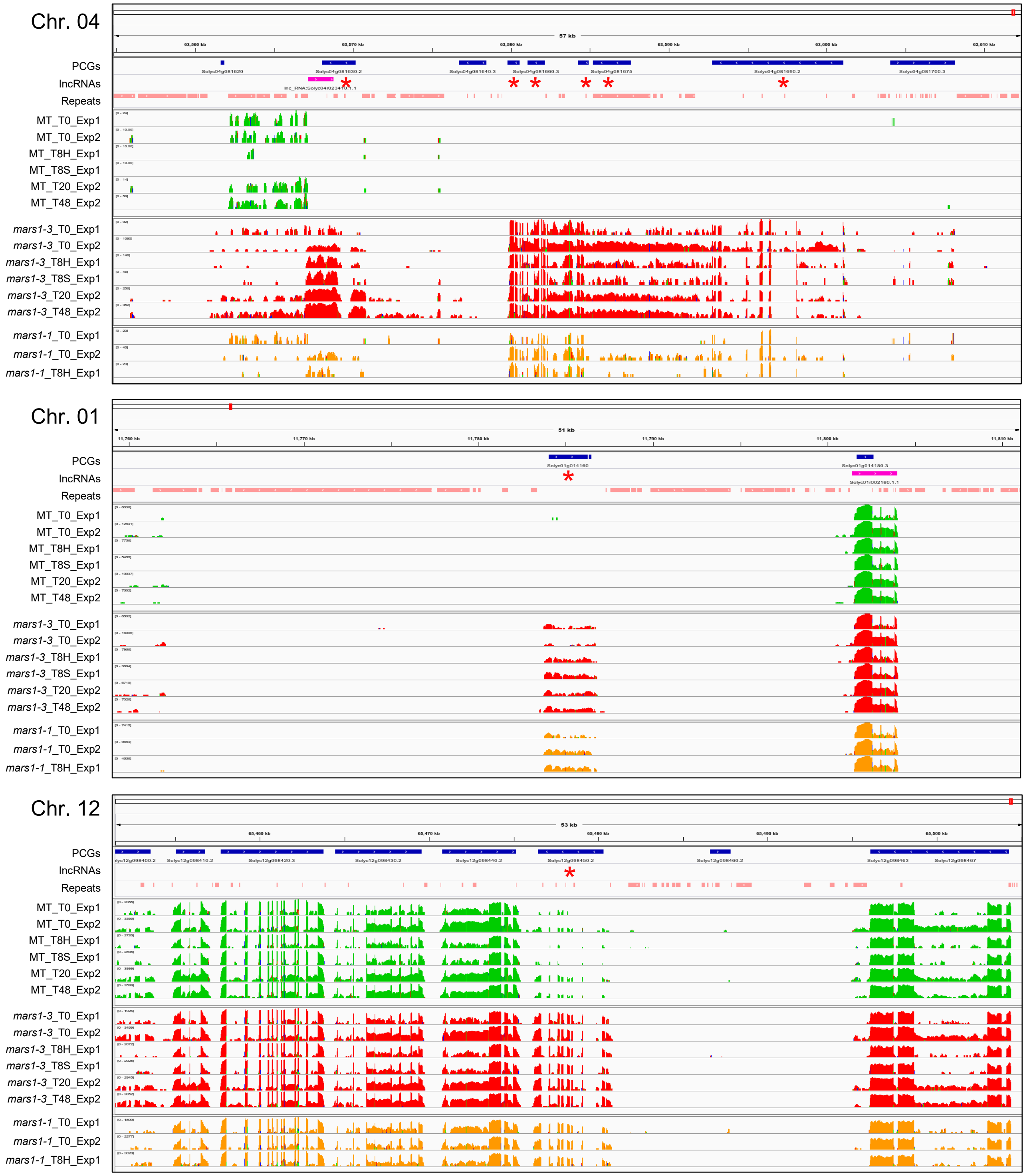


**Suppl. Figure S7. An IGV visualization of gene expression in genomic regions flanking upregulated DEGs in *mars1/rough* mutants.** The IGV visualization of RNA-seq reads in several representative regions is presented. The colored peaks indicate the coverage of reads from multiple replicates at each time point, genotype, or condition. The MT samples are depicted in green, the *mars1-3* samples in red, and the *mars1-1* samples in orange. The annotated PCGs and lncRNAs are displayed at the top of each panel in blue and purple, respectively, with arrowheads indicating the direction of transcription; repetitive regions identified by RepeatModeler are shown in beige. The red asterisks indicate genes within this region that were found to be significantly upregulated in the *mars1/rough* mutants compared to MT.

### Suppl. Figure S7 (cont.)

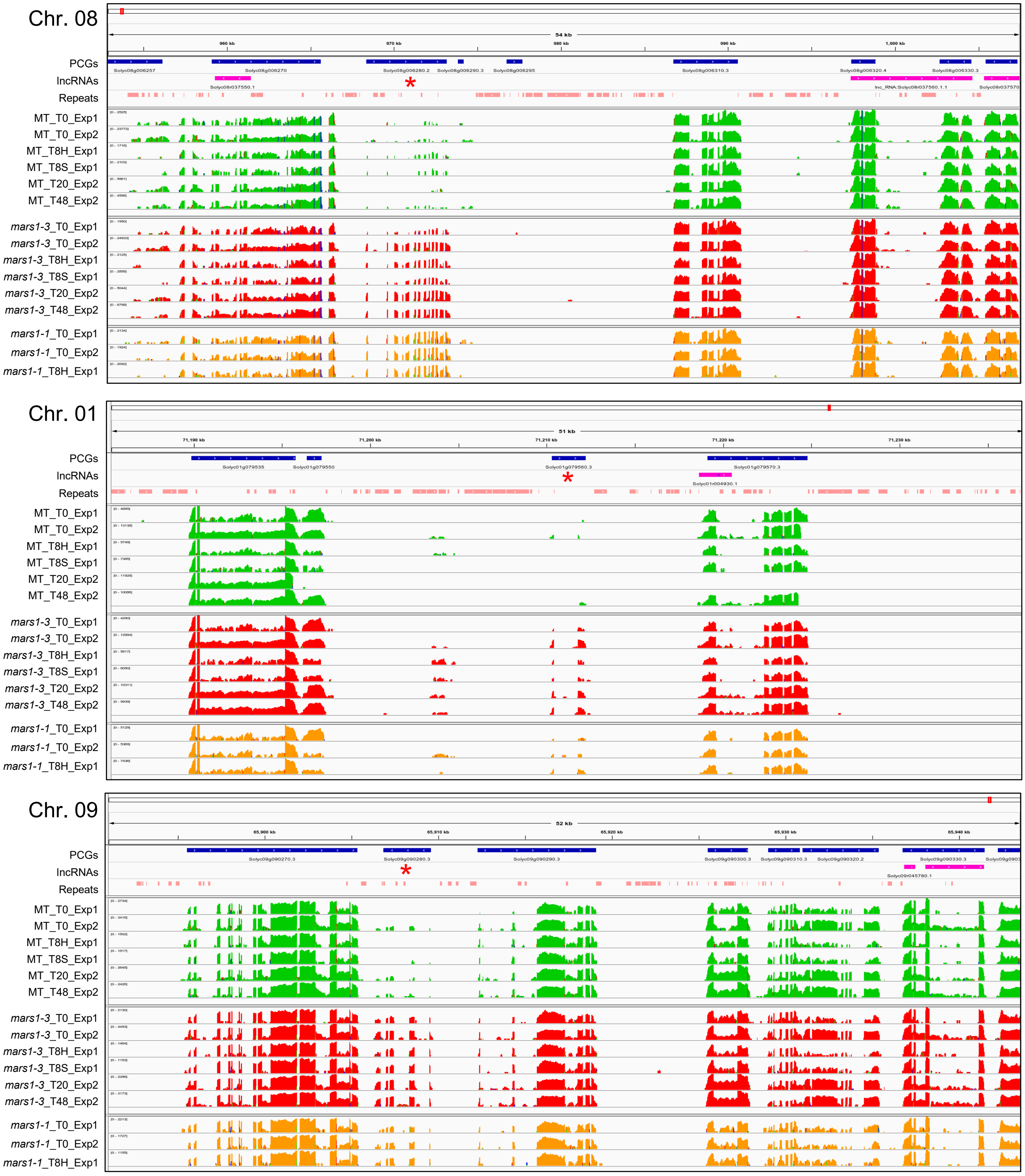


**Suppl. Figure S7. An IGV visualization of gene expression in genomic regions flanking upregulated DEGs in *mars1/rough* mutants (cont.).** The IGV visualization of RNA-seq reads in several representative regions is presented. The colored peaks indicate the coverage of reads from multiple replicates at each time point, genotype, or condition. The MT samples are depicted in green, the *mars1-3* samples in red, and the *mars1-1* samples in orange. The annotated PCGs and lncRNAs are displayed at the top of each panel in blue and purple, respectively, with arrowheads indicating the direction of transcription; repetitive regions identified by RepeatModeler are shown in beige. The red asterisks indicate genes within this region that were found to be significantly upregulated in the *mars1/rough* mutants compared to MT.

### Suppl. Figure S7 (cont.)

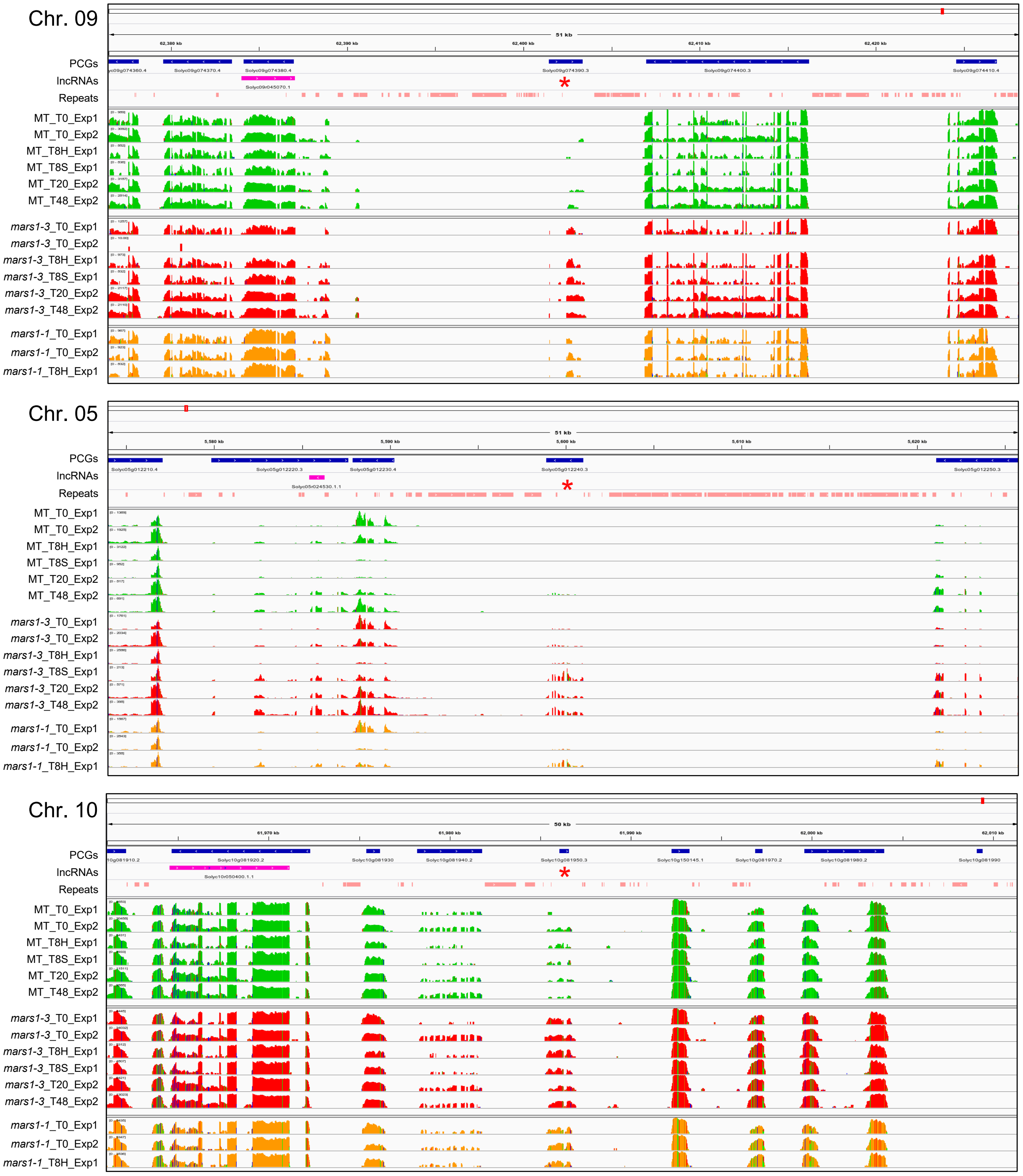


**Suppl. Figure S7. An IGV visualization of gene expression in genomic regions flanking upregulated DEGs in *mars1/rough* mutants (cont.).** The IGV visualization of RNA-seq reads in several representative regions is presented. The colored peaks indicate the coverage of reads from multiple replicates at each time point, genotype, or condition. The MT samples are depicted in green, the *mars1-3* samples in red, and the *mars1-1* samples in orange. The annotated PCGs and lncRNAs are displayed at the top of each panel in blue and purple, respectively, with arrowheads indicating the direction of transcription; repetitive regions identified by RepeatModeler are shown in beige. The red asterisks indicate genes within this region that were found to be significantly upregulated in the *mars1/rough* mutants compared to MT.

### Suppl. Figure S7 (cont.)

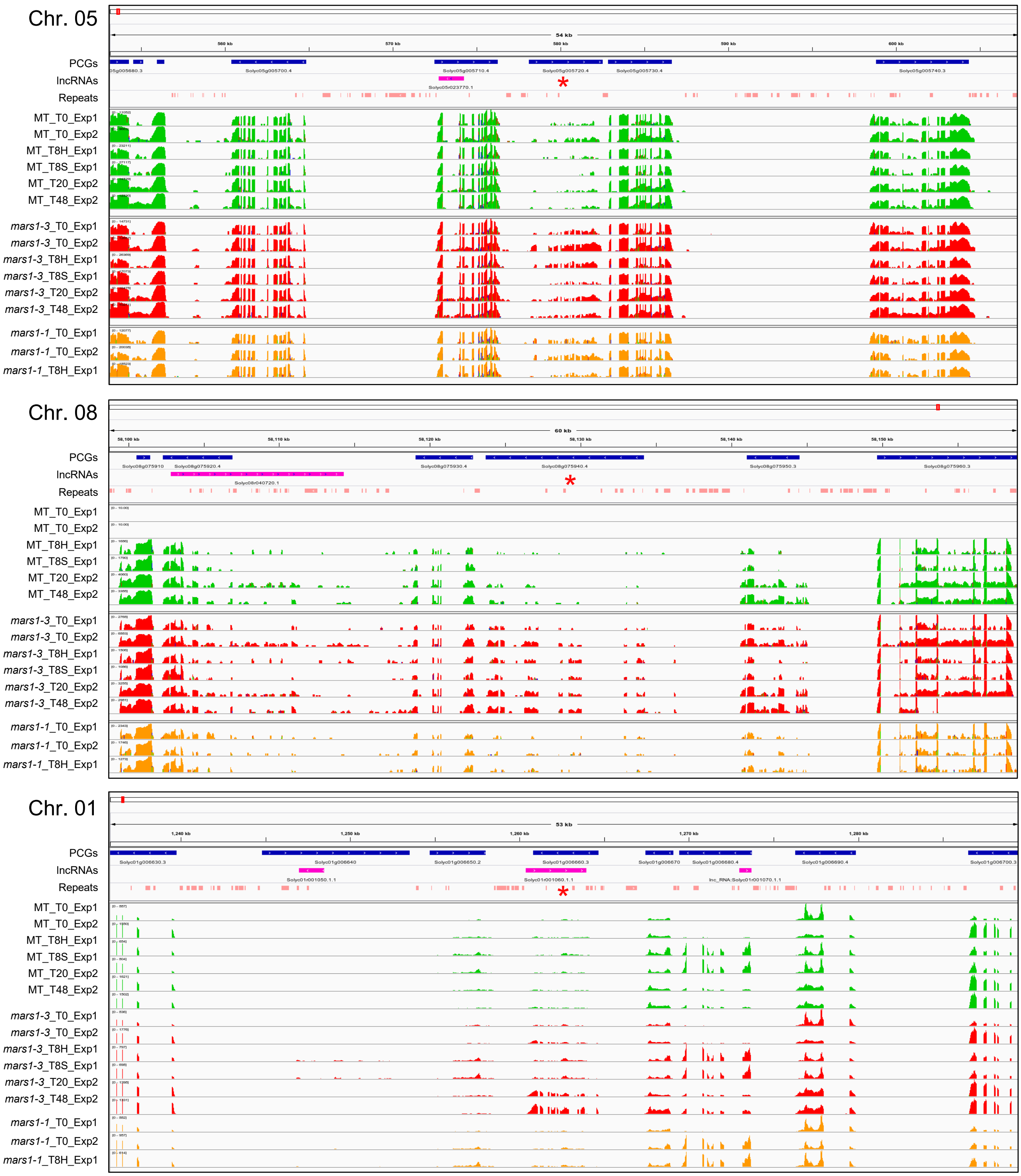


**Suppl. Figure S7. An IGV visualization of gene expression in genomic regions flanking upregulated DEGs in *mars1/rough* mutants (cont.).** The IGV visualization of RNA-seq reads in several representative regions is presented. The colored peaks indicate the coverage of reads from multiple replicates at each time point, genotype, or condition. The MT samples are depicted in green, the *mars1-3* samples in red, and the *mars1-1* samples in orange. The annotated PCGs and lncRNAs are displayed at the top of each panel in blue and purple, respectively, with arrowheads indicating the direction of transcription; repetitive regions identified by RepeatModeler are shown in beige. The red asterisks indicate genes within this region that were found to be significantly upregulated in the *mars1/rough* mutants compared to MT.

### Suppl. Figure S8

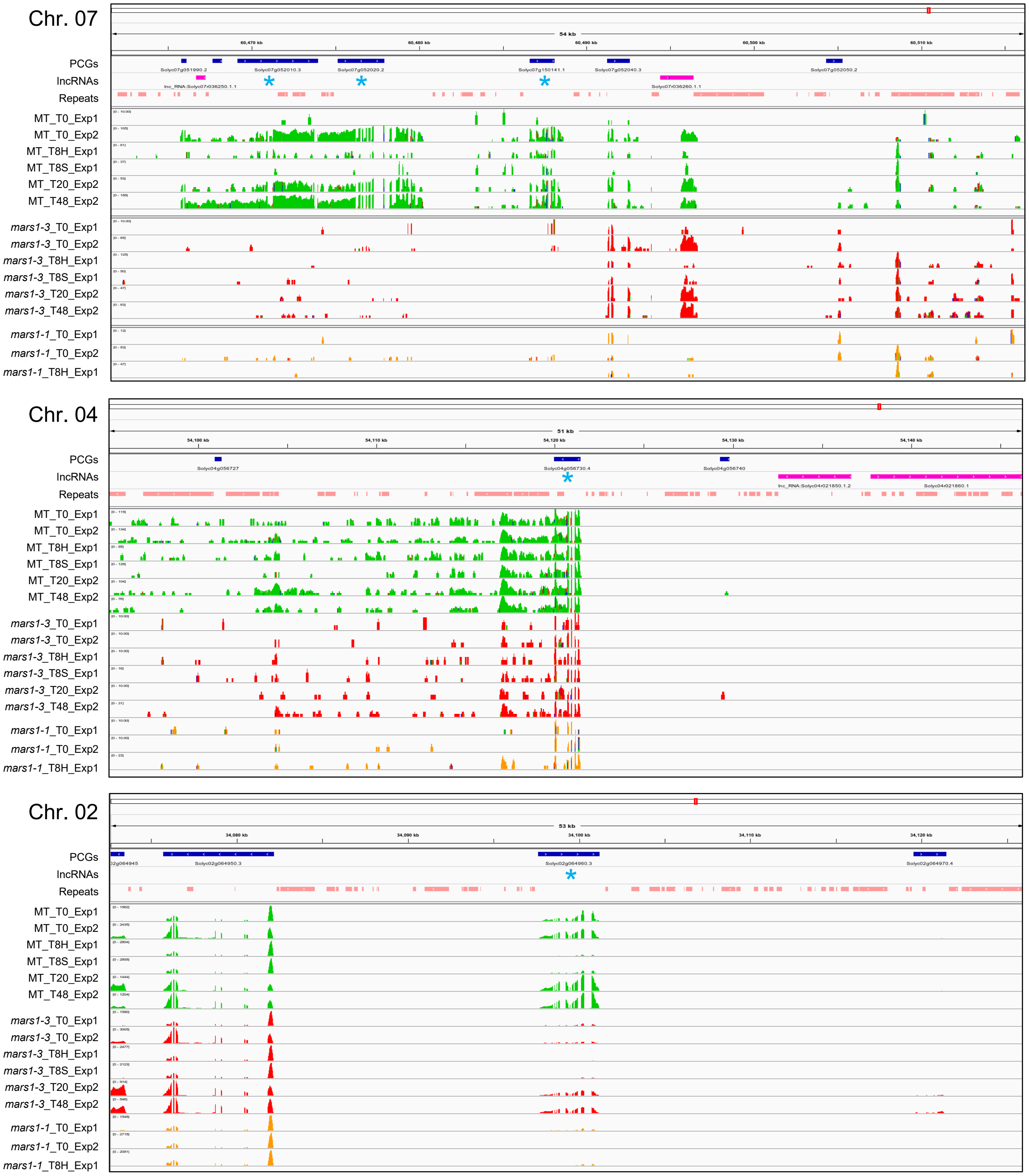


**Suppl. Figure S8. An IGV visualization of gene expression in genomic regions flanking downregulated DEGs in *mars1/rough* mutants.** The IGV visualization of RNA-seq reads in several representative regions is presented. The colored peaks indicate the coverage of reads from multiple replicates at each time point, genotype, or condition. The MT samples are depicted in green, the *mars1-3* samples in red, and the *mars1-1* samples in orange. The annotated PCGs and lncRNAs are displayed at the top of each panel in blue and purple, respectively, with arrowheads indicating the direction of transcription; repetitive regions identified by RepeatModeler are shown in beige. The blue asterisks indicate genes within this region that were found to be significantly downregulated in the *mars1/rough* mutants compared to MT.

### Suppl. Figure S9

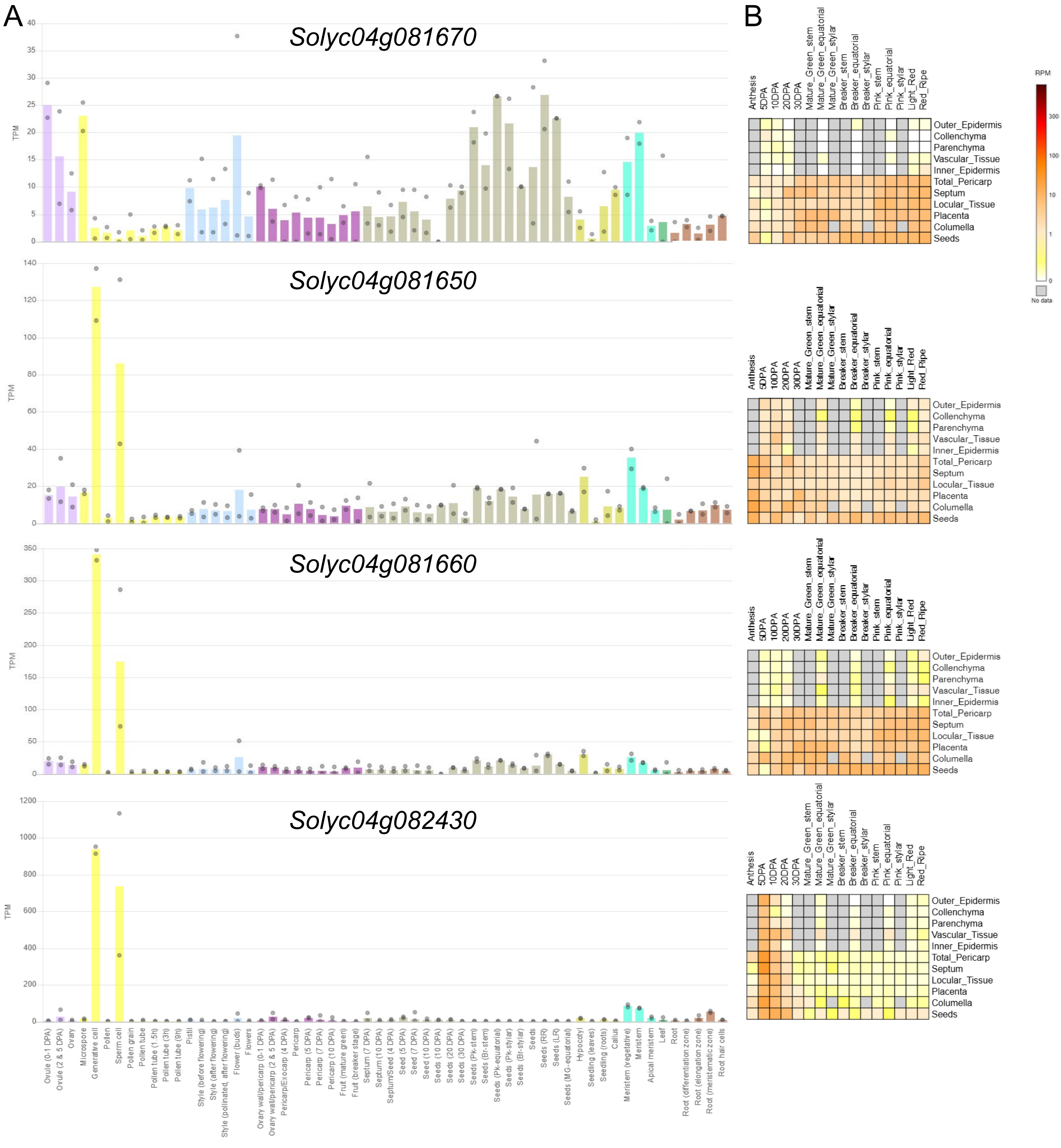


**Suppl. Figure S9. Expression profile of *Solyc04g081650*, *Solyc04g081660*, and *Solyc04g081670* in different tissues and across developmental stages.** (A) Expression values retrieved from the CoNekT RNA-seq database are given in transcripts per kilobase million (TPM), normalized for read count and gene length. Bars represent mean value; circles represent minimum and maximum values. (B) Expression data from several fruit development stages and tissues in ‘M82’ retrieved from the Tomato Expression Atlas are given in a color-coded heatmap based on reads per million mapped reads (RPM). The expression of the *Solyc04g082430* gene encoding the CycB2;4 is shown as a reference.

### Suppl. Figure S10

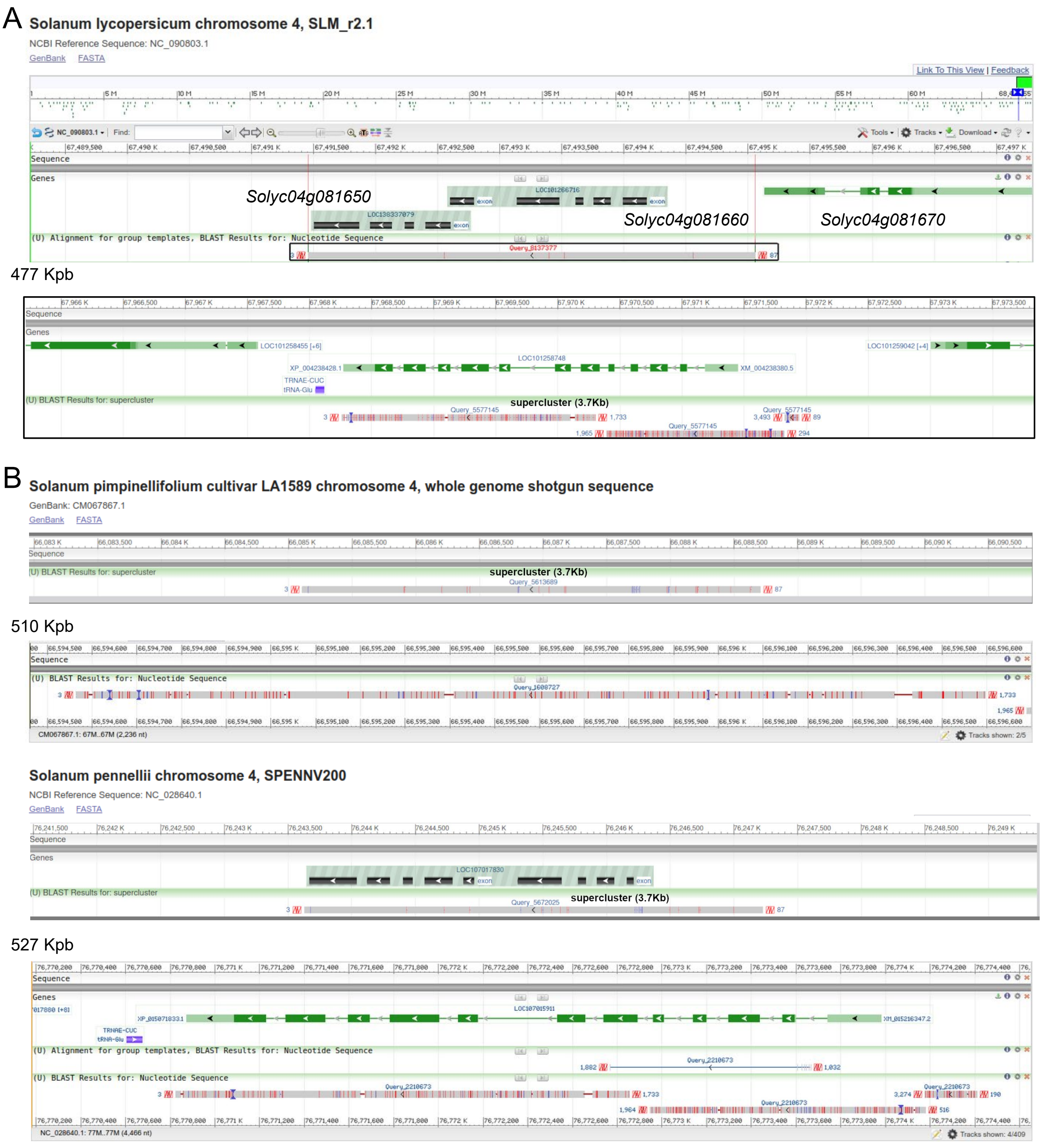


**Suppl. Figure S10. Alignment of the *Solyc04g081660-Solyc04g081650* genomic region of the *mars1/rough* mutants to the tomato genome**. (A) Visualization of the BLAST alignment against the 'Micro-Tom' reference genome (SLM_r2.1). (B) Visualization of the BLAST alignment against the *Solanum pimpinellifollium* reference genome. (C) Visualization of the BLAST alignment against the *S. pennellii* reference genome (SPENNV200). SNP polymorphisms found between the sequenced region and the reference genomes are shown as vertical red lines, and InDels are shown as horizontal red lines. The numbers on the left side of the figure indicate the distance in kbp between the two regions examined.
